# FET fusion oncoproteins disrupt physiologic DNA repair and create a targetable opportunity for ATR inhibitor therapy

**DOI:** 10.1101/2023.04.30.538578

**Authors:** Shruti Menon, Daniel Gracilla, Marcus R. Breese, Yone Phar Lin, Filemon Dela Cruz, Tamar Feinberg, Elisa de Stanchina, Ana-Florina Galic, Hannah Allegakoen, Shruthi Perati, Nicholas Wen, Ann Heslin, Max A. Horlbeck, Jonathan Weissman, E. Alejandro Sweet-Cordero, Trever G. Bivona, Asmin Tulpule

**Author notes:** Correspondence to: Asmin Tulpule MD PhD. Denotes equal contribution (Shruti Menon, Daniel Gracilla).

## Abstract

In cancers with genetic loss of specific DNA damage response (DDR) genes (i.e., BRCA1/2 tumor suppressor mutations), synthetic lethal targeting of compensatory DDR pathways has translated into clinical benefit for patients. Whether and how growth-promoting oncogenes might also create tumor-specific vulnerabilities within DDR networks is not well understood. Here we focus on Ewing sarcoma, a FET fusion oncoprotein (EWSR1-FLI1) driven pediatric bone tumor, as a model for the class of FET rearranged cancers. Native FET family members are among the earliest factors recruited to DNA double-strand breaks (DSBs), though the function of both native FET proteins and FET fusion oncoproteins in DNA repair remains to be defined. We discover that EWSR1-FLI1 and other FET fusion oncoproteins are recruited to DNA DSBs and impair the activation and downstream signaling of the DNA damage sensor ATM. In multiple FET rearranged cancers, we establish the compensatory ATR signaling axis as a collateral dependency and therapeutic target using patient-derived xenograft models. In summary, we describe how oncogenes can disrupt physiologic DNA repair and provide the preclinical rationale for specifically testing ATR inhibitors in FET rearranged cancers as part of ongoing early phase clinical trials.

---

The DNA damage response (DDR) is a tightly regulated and redundant network that tailors specific repair complexes to address a diverse set of genotoxic insults^1^. In cancer, oncogene-induced hyper-proliferation and replication stress activate the DDR which can trigger senescence or cell death programs^2,3^. Thus, the physiologic DDR represents an important barrier to oncogenic transformation and genetic loss of tumor-suppressive DDR pathways or downstream effectors are frequent cooperating events in cancer (e.g., ATM or p53 mutations)^4,5^. Loss of these DDR genes can also create collateral dependencies and therapeutic opportunities in specific cancers as exemplified by the use of poly-ADP ribose polymerase (PARP) inhibitors to target defective homologous recombination (HR) repair in BRCA1/2 mutant breast and ovarian cancers^6,7^. Outside of HR deficient cancers, while multiple synthetic lethal DDR dependencies have been described (e.g., oncogene-induced replication stress and ATR^3^), the clinical benefit of DDR-targeting therapies is less established and the mechanisms underlying many of these aberrant DDR dependencies remain to be defined^8^. More generally, whether and how oncogenes themselves regulate physiologic DNA repair to create functional defects or dependencies within DDR networks is poorly understood.

The FET family of intrinsically disordered proteins (FUS, EWSR1, TAF15) are frequent 5’ oncogenic transcription factor (TF) fusion partners in a diversity of sarcomas and leukemias^9^. These TF fusion oncoproteins are often the sole oncogenic driver alteration in these cancers and due to the difficulty in pharmacologic targeting of TFs, precision medicine approaches have been lacking. The most studied cancer in this class is Ewing sarcoma (ES), a pediatric bone tumor driven by the EWSR1-FLI1 TF fusion oncoprotein. Patients with relapsed or metastatic ES continue to have dismal outcomes despite maximally intense combination cytotoxic chemotherapy regimens^10^. Paradoxically, decades of clinical experience and laboratory testing of ES cancer cell lines have shown ES to be among the most chemo- and radiosensitive cancers^11–14^, at least initially. It has long been hypothesized that ES tumors harbor an intrinsic DNA repair defect that explains their underlying sensitivity to DNA damaging therapies. The prevailing model is that ES belongs to a family of “BRCA-like” tumors that are functionally deficient in BRCA1 (due to sequestration of BRCA1 protein by RNA-DNA hybrid structures known as R-loops) and therefore defective in HR double-strand break (DSB) repair^15^. However, both xenograft studies^13^ and clinical trials in ES patients^16^ failed to demonstrate any benefit for PARP inhibitor monotherapy, in stark contrast to the impressive clinical responses seen across HR deficient BRCA mutant and BRCA-like cancers^17,18^. Thus, the precise nature of the DNA repair defect in ES remains uncertain.

Native FET family proteins contain an N-terminal intrinsically disordered region (IDR) required for interactions amongst FET family members and a C-terminal domain with positively charged RGG (arginine-glycine-glycine) repeats, which mediate recruitment to DSBs via high affinity interactions with negatively charged poly-ADP ribose (PAR) molecules^19,20^. All 3 FET family members are rapidly recruited to DSBs in a PARP-dependent manner^20,21^, where they undergo liquid-liquid phase separation that is thought to enable compartmentalization of DSB repair proteins^22–24^, though the specific role of FET proteins in DSB repair is not well defined. Interestingly, all oncogenic FET fusion proteins including EWSR1-FLI1 share a similar structure: the N-terminal IDR of the FET protein fused to the DNA binding domain of a transcription factor (e.g., FLI1), with loss of the C-terminal RGG repeats^25^.

In this study, we address the role of the oncogenic fusion protein EWSR1-FLI1 in regulating the DNA damage response in ES. Contrary to the current classification of ES as a BRCA-like tumor with defective HR^15^, we nominate suppression of ATM activation and signaling as a new DDR lesion in ES cells and identify the compensatory ATR signaling axis as a synthetic lethal therapeutic strategy for the class of FET rearranged cancers.

## Results

### ES cells are dependent on specific HR factors for survival

To address the uncertainty surrounding putative DNA repair defect(s) in ES, we set out to identify genes that modulate ES cell survival in response to doxorubicin, a DNA DSB-inducing agent and major component of current chemotherapy regimens^26^ (Fig. 1A). We selected a CRISPR interference (CRISPRi) based screening approach given the concern of studying DSB repair phenotypes using an active Cas9 that generates DSBs^27,28^. Surprisingly, we found that key HR factors BRCA1 and PALB2 were essential for the growth of ES cells even in the absence of doxorubicin (Fig. 1B). Given the prevailing model that ES is a functionally HR-deficient, BRCA-like tumor^15^, the screen results were unexpected. We validated the finding of specific HR factor dependency using two independent guide RNAs (gRNAs) against BRCA1 and PALB2 in the screening cell line (A673) and two additional ES cell lines TC-252 and ES-8 (Figs. 1C-E and S1A, B). The same BRCA1 and PALB2 gRNAs had limited effects on cell growth in two non-ES cancer cell lines, which was consistent with the set of published CRISPRi screens^27,29,30^ and suggests the observed dependency on HR factors may be specific to ES cells (Figs. S1C, D). To test the role of EWSR1-FLI1 in inducing this dependency, we utilized an ES cell line with a doxycycline-inducible shRNA against EWSR1-FLI1^31^ (Fig. S1E). EWSR1-FLI1 knockdown rescued the growth defects caused by BRCA1 or PALB2 loss, confirming that the oncogenic fusion protein is necessary for the observed dependency on these specific HR factors in ES cells (Fig. 1F).

**Figure 1:**
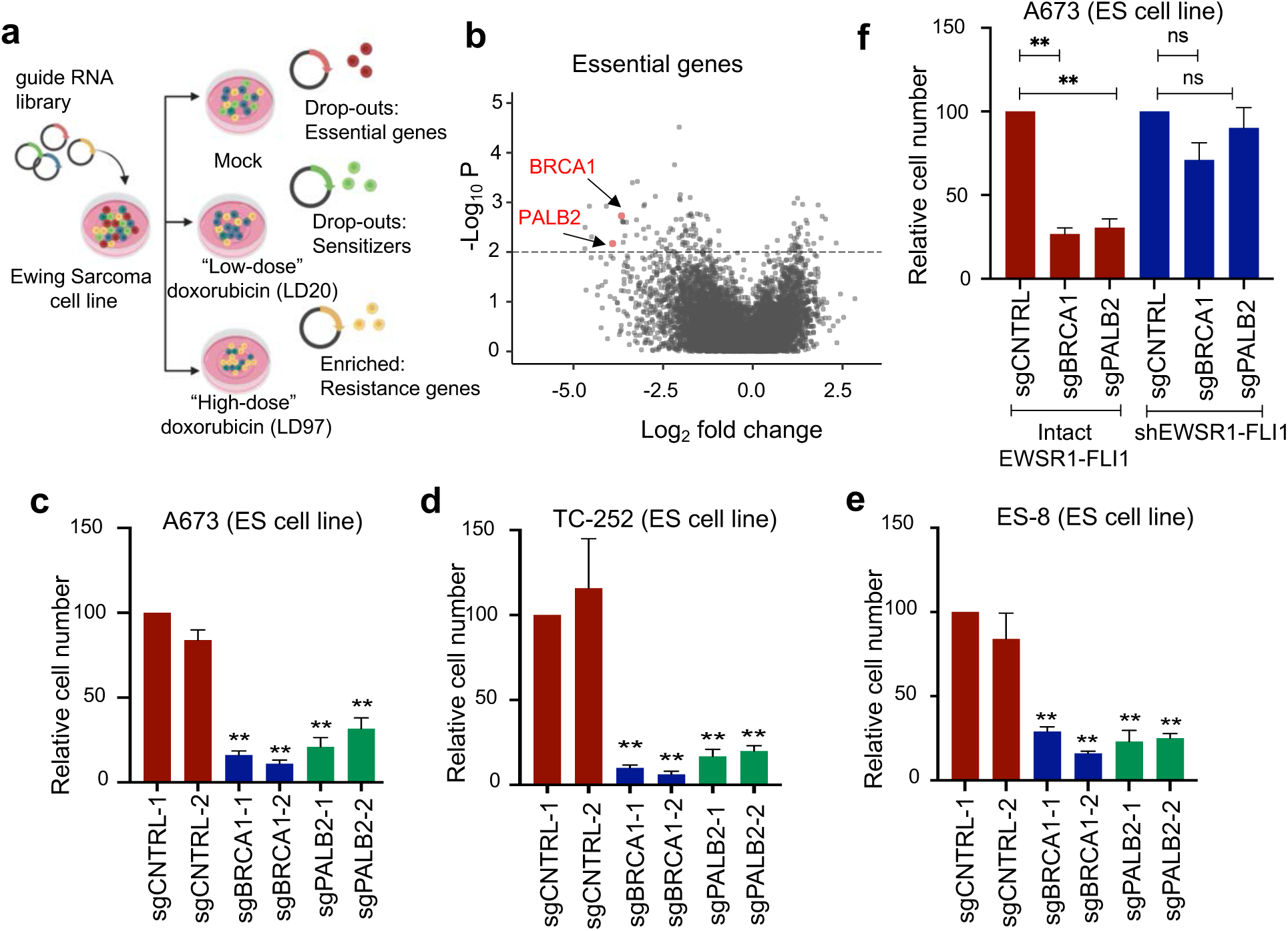
ES cells are dependent on specific HR factors for survival. A) Schematic of CRISPRi genetic screen of 7,000 cancer associated genes in ES cell line A673. B) Volcano plot of sgRNA log_2_fold change (X-axis) vs -log10(p value), average of 2 biologic replicates. Full screen results in Supplementary Table 1. C-E) Growth assays in dCas9 expressing ES cell lines upon introduction of 2 independent sgRNAs against BRCA1, PALB2, or control. n = 4. F) Growth assays in ES cell line A673 with doxycycline-inducible shRNA against EWSR1-FLI1. For all panels, error bars represent ± SEM, ^∗∗^p < 0.01 by one-way ANOVA with post hoc Tukey’s HSD test.

CRISPRi screening also identified genes whose loss sensitized ES cells to doxorubicin including LIG4 and NHEJ1/XLF (Fig. S1F). The presence of key canonical non-homologous end joining (c-NHEJ) genes as top chemo-sensitizer hits validated our experimental approach as this pathway is critical for repairing both drug and ionizing radiation (IR)-induced DSBs and provides evidence for a functional c-NHEJ pathway in ES cells. The final category of screen hits were genes whose loss promoted survival under high doses of doxorubicin (LD97), mirroring the residual disease state in ES patients (Fig. S1G). SLFN11 is a notable hit as loss of SLFN11 has been shown to promote chemotherapy resistance in multiple cancer subtypes including ES^32,33^. The complete screen results are provided in Supplementary Table 1 (see Supplemental Note 1 for discussion of DepMap results). Given the paradoxical finding of HR factor dependence in ES, we chose to focus on the regulation of DSB repair in ES and whether ES tumors are properly classified as BRCA-like cancers.

### ES patient tumors do not display the genomic scars of HR deficiency

We next directly examined genomic DNA from 100 ES patient tumor samples^34^ for evidence of functional HR deficiency. This is a validated strategy in BRCA1/2 mutant and BRCA-like tumors wherein defective HR repair results in specific genomic scars that result from increased utilization of compensatory pathways such as alternative end-joining (alt-EJ) and single-strand annealing (SSA)^35,36^. We developed a custom bioinformatics pipeline to analyze the genomic landscape of ES tumor genomes, using BRCA1/2 mutant and wild-type breast cancer genomes (from EGAD00001001322^35^) to validate our algorithms. The ES cohort was reported in Tirode et al to represent diagnostic, pre-treatment biopsies of the primary tumor site, with 63% of patients having localized disease^34^. The ES whole genome sequencing data were previously deposited in the European Genome-phenome Archive (EGAS00001000855, EGAS00001000839).

At least 99 (single base substitution) mutational signatures have been identified in human cancers with Signature 3 being the most highly associated with BRCA1/2 mutant cancers^35^. Our analysis confirmed high levels of Signature 3 in BRCA mutant as compared to BRCA wild-type tumors but did not show increased levels of Signature 3 in ES tumors (Figs. 2A and S2A). In addition to increased Signature 3, BRCA1/2 mutant tumors displayed an increase in the number and size of deletions compared to BRCA wild-type tumors (Figs. 2B, C and S2B-D) consistent with previous reports^35^. ES tumors are clearly distinct from BRCA1/2 mutant tumors displaying few deletions per genome and a size distribution skewed further towards small deletions than even BRCA wild-type tumor samples (Figs. 2B, C and S2B-D).

**Figure 2:**
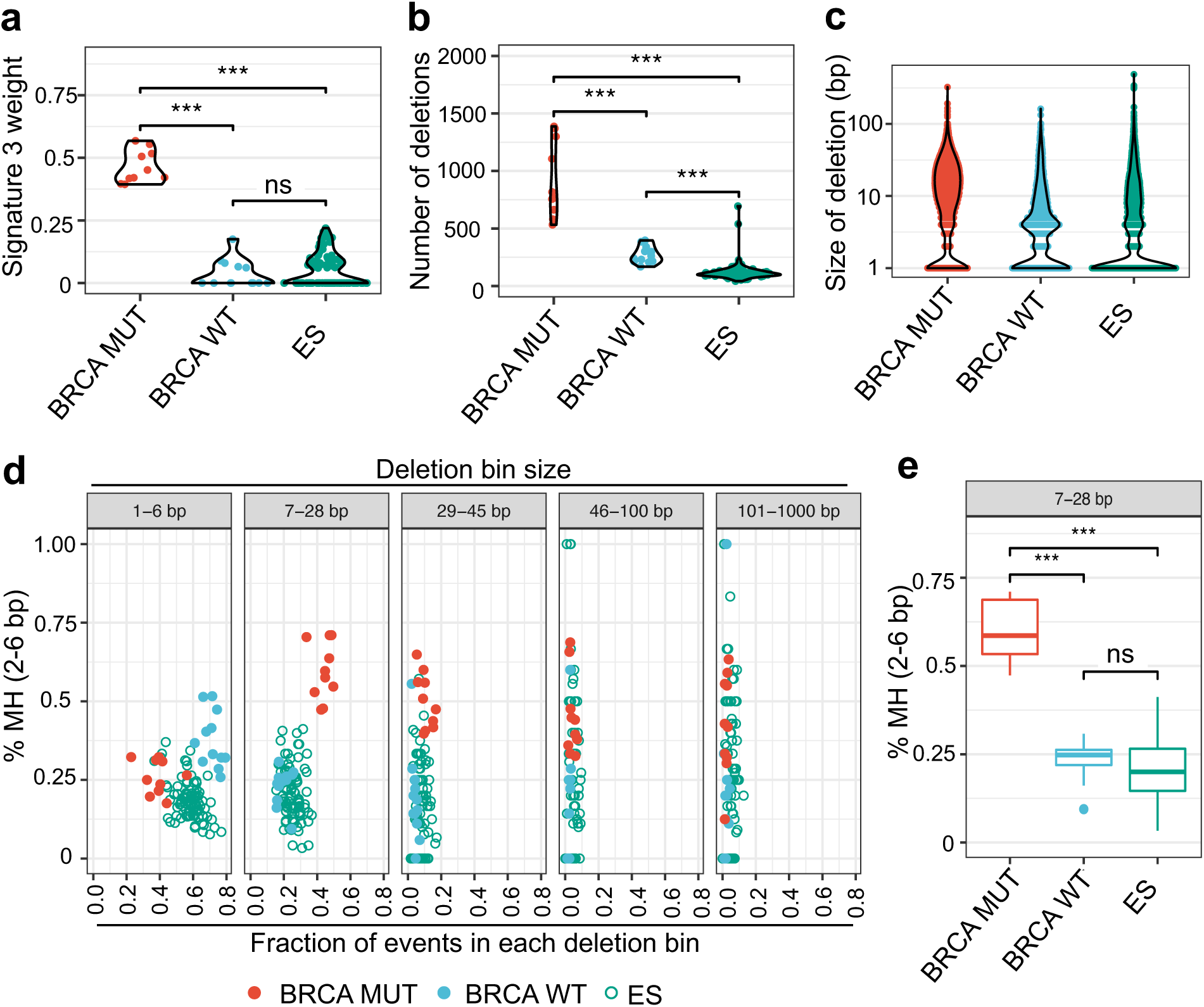
ES patient tumors do not display the genomic scars of HR deficiency. A) Weight of mutational signature 3 shown for BRCA mutant, wild-type, and ES tumor samples. B) Total number of unique deletions per sample. C) Size of MuTect deletions compared across tumor groups. D) For all samples, deletions were first subdivided into specific base pair (bp) length bins, then the fraction of deletions with 2-6bp microhomology (MH) was plotted (Y-axis) vs. the fraction of deletions present in this bin (X-axis). E) Breakout of the 7-28bp deletion length bin showing the distribution of deletions with microhomology (2-6bp). For all panels, *** denotes p< 0.001, ns denotes not significant by Mann-Whitney test. European Genome-phenome Archive accession numbers: ES samples (EGAS00001000855 - Institute Curie cohort and EGAS00001000839 - St. Jude’s cohort)^34^, breast cancer samples (EGAD00001001322)^35^.

We further examined the sequences flanking deletion sites for short stretches of overlapping microhomology (MH). Defective HR repair in BRCA mutant or BRCA-like tumors results in increased usage of the error prone DSB repair pathway alt-EJ that employs MH for initial annealing of resected DSB ends^37^. We utilized BRCA mutant/wild-type samples to computationally define deletion size bins and identified an increase in the proportion of intermediate size deletions (7-28 base pairs (bp), 29-45 bp) in BRCA mutant samples as reported previously (Figs. S2E, F). We then verified a significant increase in breakpoint MH at intermediate size (7-28bp and 29-45bp) deletions in BRCA mutant tumors compared to wild-type and no increase in MH at small deletions (1-6bp) where c-NHEJ predominates (Figs. 2D, E, and S2E-G), both consistent with published findings^35^. ES samples clustered with BRCA wild-type tumors and display a trend toward even less MH-mediated DSB repair than BRCA wild-type samples (Figs. 2D, E, and S2G). Our findings in ES were independent of the recurrent STAG2 and p53 mutations that occur in a subset of these cancers (Figs. S2H, I). Taken together, these results demonstrate that ES patient tumors do not display the genomic signatures of BRCA1/2 mutant cancers and lack the footprint of isolated HR deficiency.

### EWSR1-FLI1 reduces resection-dependent DSB repair

The absence of HR deficient genomic scars in ES tumors and paradoxical requirement of specific HR factors (e.g., BRCA1) for ES cell survival prompted us to systematically re-examine how EWSR1-FLI1 impacts DSB repair pathway utilization (Fig. 3A). We posited that previous reports of defective HR in ES might alternatively be explained by a more general upstream defect in DSB repair. We utilized a set of well-established DSB repair reporter cell lines wherein expression of the I-SceI endonuclease induces a DSB within an interrupted GFP reporter cassette, such that utilization of a particular DSB repair pathway restores the GFP coding sequence enabling a quantitative readout of individual repair pathway efficiency^38^. The use of fluorescently tagged plasmids enabled detection of DSB repair events specifically in cells that expressed both EWSR1-FLI1 (or empty vector) and I-SceI (Fig. S3A).

**Figure 3:**
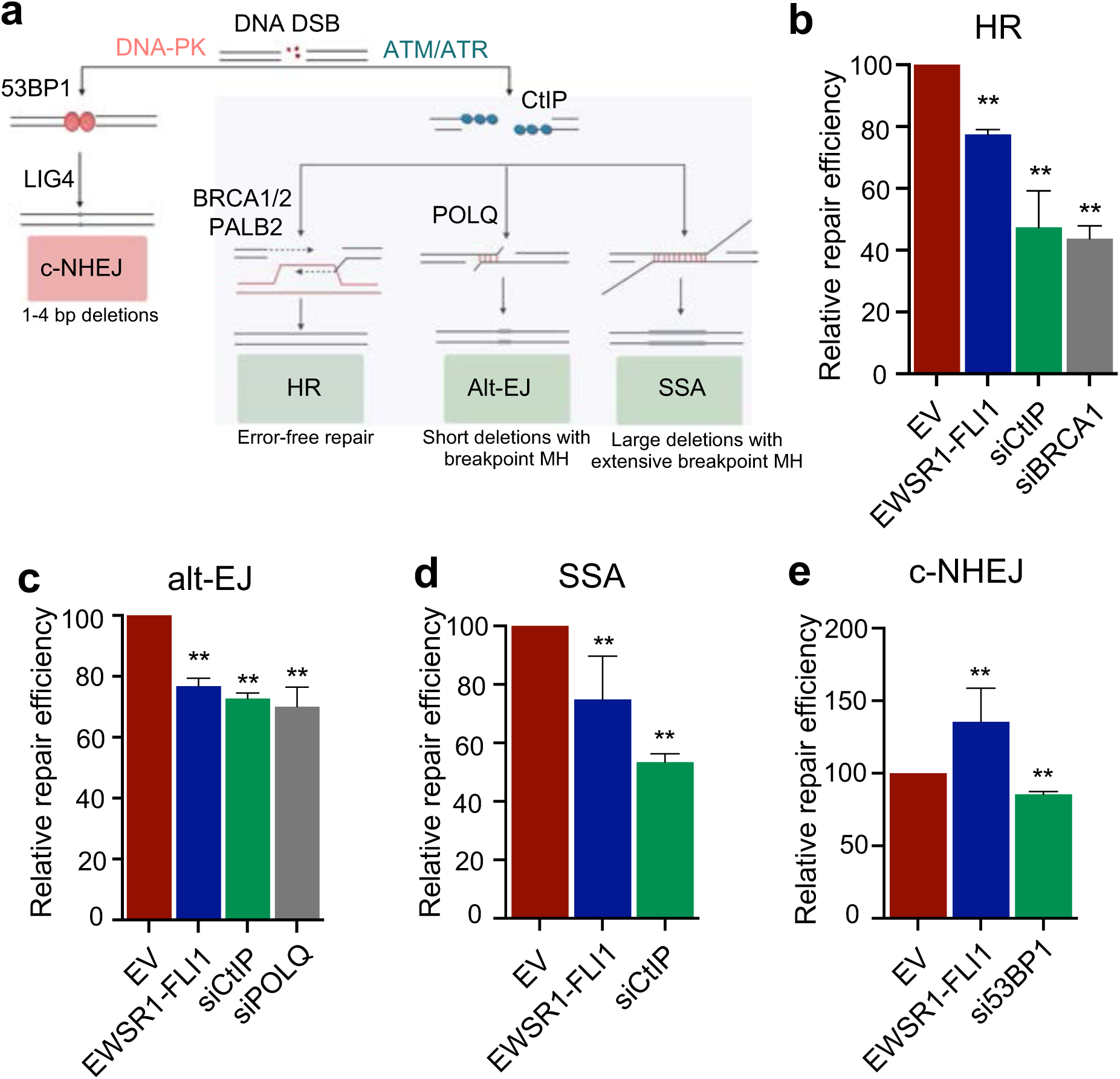
EWSR1-FLI1 reduces resection-dependent DSB repair. A) Schematic of DSB repair. DSBs either undergo direct repair via c-NHEJ or bi-directional end-resection to create an intermediate for HR, alt-EJ, or SSA. B-E) Relative repair efficiency as measured by GFP positivity in pathway-specific DSB repair reporters for HR (DR-GFP), alt-EJ (EJ2), SSA, and c-NHEJ (EJ5) upon expression of empty vector (EV), EWSR1-FLI1, or siRNA gene knockdown for 72 hours. For all panels, error bars represent ± SEM, ** denotes p < 0.01 by one-way ANOVA with post hoc Tukey’s HSD test. n=4 for all panels.

To examine HR repair, we utilized the DR-GFP reporter and observed a small reduction in HR upon EWSR1-FLI1 expression, consistent with previous reports^15^ (Fig. 3B). However, EWSR1-FLI1 expression also reduced the efficiency of MH-mediated alt-EJ repair (EJ2) and long-stretch MH mediated-SSA repair (Figs. 3C, D). EWSR1-FLI1 mediated reductions in HR, alt-EJ, and SSA repair efficiency were modest compared to knockdown of the key end-resection factor CtIP that is required for all resection-dependent DSB repair^39^ or the relevant pathway specific controls (Figs. 3A-C). These data indicate EWSR1-FLI1 expression does not result in isolated HR defects, but instead an intermediate reduction in the efficiency of all three resection-dependent DSB repair pathways.

We also evaluated c-NHEJ, a fast-acting DSB repair pathway which does not require end-resection, and observed an increase in c-NHEJ utilization upon EWSR1-FLI1 expression using the EJ5 reporter system (Fig. 3E). Using a distinct I-SceI based DSB repair reporter^40^, we directly examined the junctional sequences after DSB induction for evidence of c-NHEJ or MH-mediated alt-EJ repair (HR and SSA events are not evaluable in this system) and confirmed an increase in c-NHEJ and decrease in alt-EJ upon expression of EWSR1-FLI1 (Fig. S3B). We verified that the repair phenotypes were not the result of EWSR1-FLI altering cell cycle profiles or causing a cell cycle arrest in these short-term assays (Figs. S3C, D). In summary, we demonstrate that EWSR1-FLI1 does not induce isolated HR deficiency but instead moderately reduces the efficiency of all three resection-dependent DSB repair pathways.

### ATM activation and signaling is suppressed by the Ewing sarcoma fusion oncoprotein

To explain our finding that EWSR1-FLI1 expression leads to intermediate reductions in resection-dependent DSB repair efficiency, we focused on the upstream regulation of the DDR in ES cells (Fig. 3A). An important component of the initial DDR is a partially redundant signaling network with three principal kinases, DNA-PK, ATM, and ATR, each of which control distinct aspects of DDR signal amplification and DSB repair pathway choice^41^. We tested whether EWSR1-FLI1 affects the activation and function of these three apical DDR kinases; in particular, ATM was a logical candidate since it promotes DNA end-resection and ATM loss creates a synthetic lethal dependence on specific HR proteins such as BRCA1^42,43^.

We utilized 3 distinct ES cell line models with inducible EWSR1-FLI1 depletion: A673 and TC-32 ES cells with doxycycline-inducible shRNA’s targeting EWSR1-FLI1 (either the junctional sequence or the 3’ FLI1 portion of the fusion, see Methods) and A673 “EWSR1-FLI1 degron” cells with an auxin-inducible degron (AID) tag inserted at the endogenous EWSR1-FLI1 locus^44^. Short-term EWSR1-FLI1 depletion did not impact the baseline low-level activation of the three DDR kinases or cell viability in the absence of exogenous DNA damage (auxin-mediated depletion is within 3 hours and maximal shRNA depletion is ∼70% at 72 hours) (Figs. 4A-D, and S4A-F). However, depletion of EWSR1-FLI1 prior to ionizing radiation (IR) treatment increased both ATM activation (autophosphorylation) and downstream ATM signaling (phosphorylation of key downstream targets CHK2 and KAP1) consistently across all three ES cell systems (Figs. 4A-D and S4A, B). In contrast, IR or radiomimetic neocarzinostatin (NCS)-dependent activation of the other apical DDR kinases (DNA-PK, ATR) was unaffected by EWSR1-FLI1 knockdown in ES cells (Figs. S4C-S4F).

**Figure 4:**
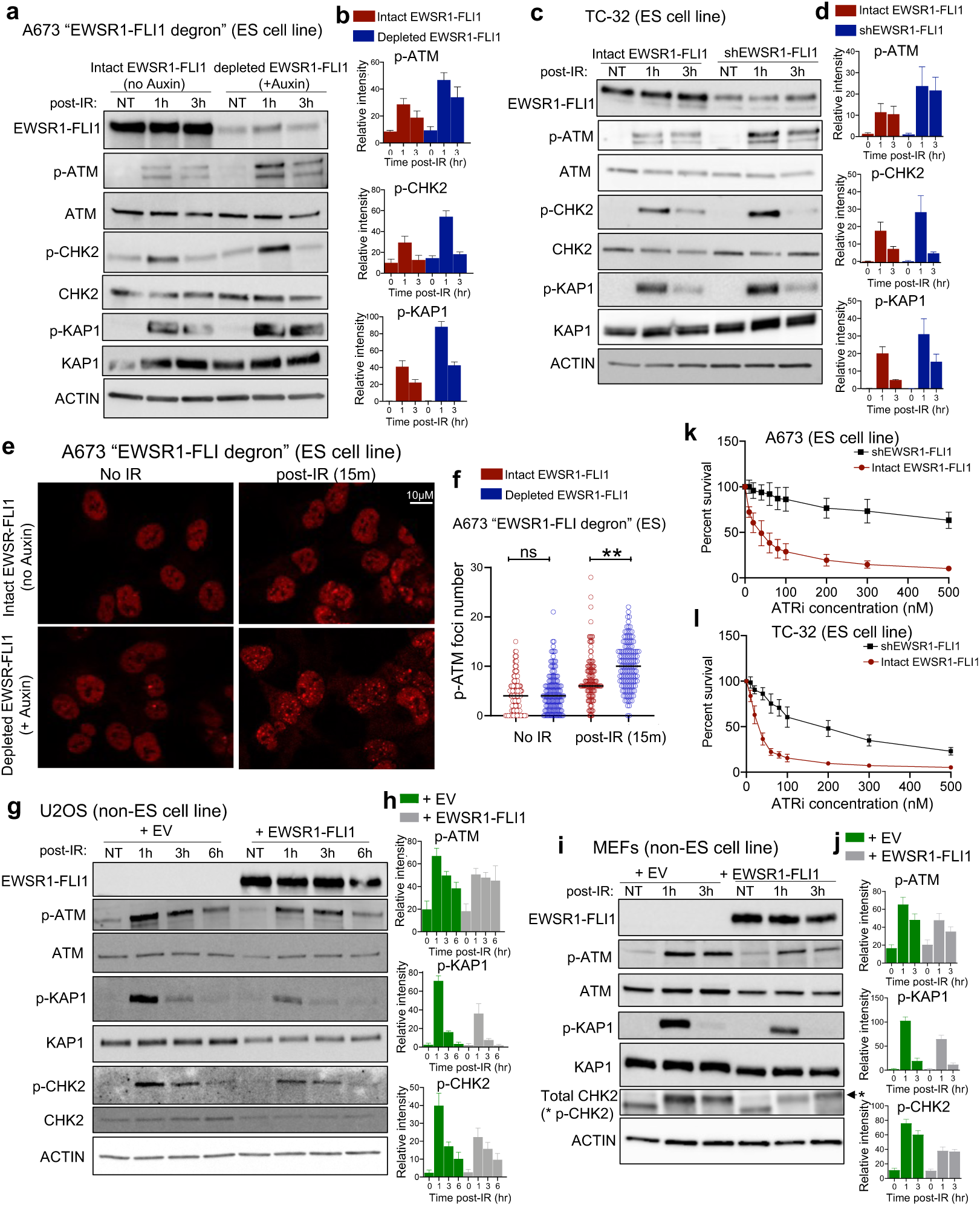
ATM activation and signaling is suppressed by EWSR1-FLI1. A-D) Western blotting and quantification upon IR (5Gy) in A673 “EWSR1-FLI1 degron” ES cells +/- 200 μM IAA (auxin) for 3 hours to degrade endogenous EWSR1-FLI1 prior to IR or TC-32 ES cells with doxycycline-inducible shRNA against EWSR1-FLI. E, F) Representative images and quantification of p-ATM foci number in A673 “EWSR1-FLI1 degron” ES cells +/- 200 μM IAA for 3 hours to deplete EWSR1-FLI1 prior to 0.5 Gy IR. ** denotes p < 0.01 by unpaired t-test. G-J) Representative Western blots and quantification upon IR (5 Gy) at indicated time points in non-ES cell lines (U2OS, MEFs) upon expression of EV (empty vector) or EWSR1-FLI1. K, L) ATR inhibitor (elimusertib) dose-response curves for ES cell lines A673 and TC-32 with dox-inducible shRNA against EWSR1-FLI1, p<0.05 for both comparisons, paired t-test. For all panels, error bars represent ± SEM for at least 4 replicates for each panel.

We measured p-ATM foci formation in A673 “EWSR1-FLI1 degron” ES cells as a complementary test for ATM activation. In the absence of exogenous DNA damage, depletion of EWSR1-FLI1 did not alter p-ATM foci number (Figs. 4E, F). Upon IR treatment, ES cells with intact endogenous EWSR1-FLI1 display a minimal increase in p-ATM foci, whereas rapid depletion of EWSR1-FLI1 prior to IR significantly increased p-ATM foci number, consistent with our Western blotting results (Figs. 4E, F). The rapid (3 hour) depletion of EWSR1-FLI1 and unchanged basal levels of DDR kinase activation suggest the observed ATM effects are not secondary downstream consequences from loss of the oncogenic driver in ES cells, but instead may result from EWSR1-FLI’s partial suppression of IR-dependent ATM activation.

To test this directly, we performed the reciprocal experiment by expressing EWSR1-FLI1 in 2 non-ES cell lines (U2OS cancer cells and mouse embryonic fibroblasts, MEFs) to determine whether ectopic EWSR1-FLI1 expression could impair IR-dependent ATM activation. In both cell lines, EWSR1-FLI1 reduced ATM activation (p-ATM) and downstream ATM signaling (p-KAP1/p-CHK2) only in response to IR (and not at baseline) (Figs. 4G-J, see Methods on measuring CHK2 phosphorylation by mobility shift in MEFs^45,46^). Consistent with our finding in ES cells, ectopic EWSR1-FLI1 expression did not impact DNA-PK or ATR activation and signaling (Figs. S4G, H). These data indicate EWSR1-FLI1 (either directly or indirectly) regulates the activation and downstream signaling of ATM, without affecting the other two apical DDR kinases.

We note that EWSR1-FLI1 does not completely suppress ATM functionality, as all three modifications (p-ATM/p-CHK2/p-KAP1) are induced to some extent upon irradiation of ES cell lines. To determine the biologic and therapeutic significance of partial ATM suppression in ES cells, we performed the following studies:

1)We benchmarked our effect size to loss of well-characterized activators of ATM including the MRN (MRE11/RAD50/NBS1) complex, MDC1, RNF8 and RNF168^41^, in order to contextualize the extent of ATM suppression in ES and establish the biological significance. There is a hierarchy amongst genes regulating ATM activity in response to DSBs: the MRN complex is required for ATM activation^47,48^ and serves as an upper bound for effect size, while the scaffolding protein MDC1^49,50^ and E3 ubiquitin ligases RNF8^51–53^ and RNF168^54,55^ have an intermediate phenotype given their important but not essential role in amplifying ATM signaling. Importantly, even loss of intermediate phenotype genes (i.e., MDC1^49^, RNF8^51^ or RNF168^54^) still produces a significant functional impact in terms of increased sensitivity to DSB-inducing drugs and IR. Using both ectopic expression in MEFs and endogenous depletion of EWSR1-FLI1 in ES cells, we show the suppressive effects of EWSR1-FLI1 on ATM activation/signaling is indeed not equivalent to the near complete functional ATM deficiency seen with loss of the MRN complex (∼30-60%), but overall comparable with loss of MDC1, RNF8, or RNF168. We provide a detailed description of these experiments in Supplemental Note 2 and accompanying Supplemental Figure 7. In summary, by benchmarking EWSR1-FLI1 to loss of established ATM regulators and showing similar effect sizes, these data provide context for the magnitude of ATM suppression in ES reported here and establish the biological significance.

2) Next, we tested the functional relevance and therapeutic consequence of the partial ATM defects in ES cells. Both ATM and ATR coordinate aspects of resection-dependent DSB repair (Fig. 3A) and ATM mutant tumors display synthetic lethality with ATR inhibitors, reflecting the vital compensatory role of the ATR signaling axis in the absence of ATM^56,57^ (note ATR signaling itself is not increased in the setting of ATM loss, consistent with our findings in ES cells). We therefore asked whether the extent of ATM suppression in ES was sufficient to induce a collateral dependence on the ATR signaling axis. Indeed, A673 and TC-32 ES cells displayed nanomolar range sensitivity to the ATR inhibitor elimusertib (IC50’s of 28 and 34nM respectively, comparable to cancer cell lines with known pathogenic ATM loss of function mutations^58^), and ATR inhibitor sensitivity was significantly reversed by EWSR1-FLI1 knockdown in both cell lines (Figs. 4K, L).

Previously described mechanisms of ATR sensitivity, most notably replication stress (RS)^59^, could also contribute to the elimusertib effects that we observed. To test this directly, we expressed RNAseH1, which degrades R-loops, the major source of oncogene-induced replication stress in ES^15^, in A673 ES cells. RNAseH1 expression did not alter the ATR inhibitor sensitivity of ES cells, (Figs. S4I, J), suggesting that the molecular basis of ATR inhibitor response in ES cells may largely be a consequence of ATM suppression. EWSR1-FLI1 knockdown also reversed ES cell sensitivity to an inhibitor of ATR’s key downstream target, CHK1, and to treatment with the DSB-inducing chemotherapeutic agent doxorubicin (Figs. S4K-M), further establishing the functional relevance and therapeutic consequence of the partial ATM defects in ES cells.

We note here that the above data do not exclude additional roles for EWSR1-FLI1 in regulating the DDR that may contribute to the chemosensitivity of these tumors, and we address the genomic instability of ATM mutant tumors and the need to define mutational/deletion signatures as a comparator for ES tumors in our Discussion. In summary, our collective findings nominate partial suppression of ATM activation/signaling and resultant synthetic lethality with the compensatory ATR signaling axis as a new and therapeutically relevant DNA repair lesion in ES.

### EWSR1-FLI1 and other FET fusion oncoproteins are recruited to DNA double-strand breaks

How does EWSR1-FLI1, a transcription factor fusion oncoprotein, regulate DNA repair to suppress IR-dependent ATM activation? To assess how EWSR1-FLI1 regulates the transcription of DDR and DSB repair genes, we analyzed published RNA-sequencing data in ES cells before and after EWSR1-FLI1 knockdown^60^ (Fig. S5A). Interestingly, we found that EWSR1-FLI1 either increased or had minimal effect on the expression of key genes involved in ATM activation (e.g., MRN complex, MDC1, RNF8, RNF168, ATM, CHK2). We also readily detected these same ATM regulators by Western blotting in A673 and multiple other ES cell lines. These data suggest EWSR1-FLI1’s transcriptional effects do not create a state of low/absent DNA repair factors (e.g., MRN complex deficiency) that would explain the suppression of IR-dependent ATM activation/signaling that we observe here.

As an alternate hypothesis, we tested whether previously reported interactions between EWSR1-FLI1 and native FET proteins (e.g., EWSR1)^61,62^ might contribute to the observed ATM effects, given that native FET proteins localize to DSBs and are thought to nucleate initial compartmentalization of DSB repair factors. Consistent with these prior studies, EWSR1-FLI1 and another native FET family member (FUS) coprecipitated with FLAG-tagged EWSR1 (Fig. S5B). The strength of these interactions was unchanged in the setting of the DNA DSB-inducing agent NCS indicating a high affinity interaction not disrupted by the DDR.

We then directly tested whether the native EWSR1::EWSR1-FLI1 interaction could promote localization of the fusion oncoprotein to DNA DSBs. Indeed, we discovered that EWSR1-FLI1, like native EWSR1, is recruited to laser-induced DSBs (Figs. 5B-E). The kinetics of EWSR1-FLI1’s DSB recruitment were delayed compared to native EWSR1, whose rapid recruitment is attributed to C-terminal, positively charged arginine-glycine-glycine (RGG)-rich domains not present in EWSR1-FLI1 (Figs. 5A-E). To test how loss of these RGG domains impact DSB recruitment of EWSR1-FLI1, we reintroduced either 1 or all 3 RGG-rich domains (the entire EWSR1 C-terminus) into the fusion oncoprotein. In an RGG dose-dependent manner, the RGG containing versions of EWSR1-FLI1 displayed earlier DSB recruitment kinetics and higher levels of overall recruitment when compared to EWSR1-FLI1 (Figs. S5C, D). Moreover, the full (3) RGG domain containing version of EWSR1-FLI1 further suppressed ATM signaling beyond the effects seen with EWSR1-FLI1 (Figs. S5E, F), suggesting a potential correlation between the extent of EWSR1-FLI1 DSB recruitment and ATM suppression.

**Figure 5:**
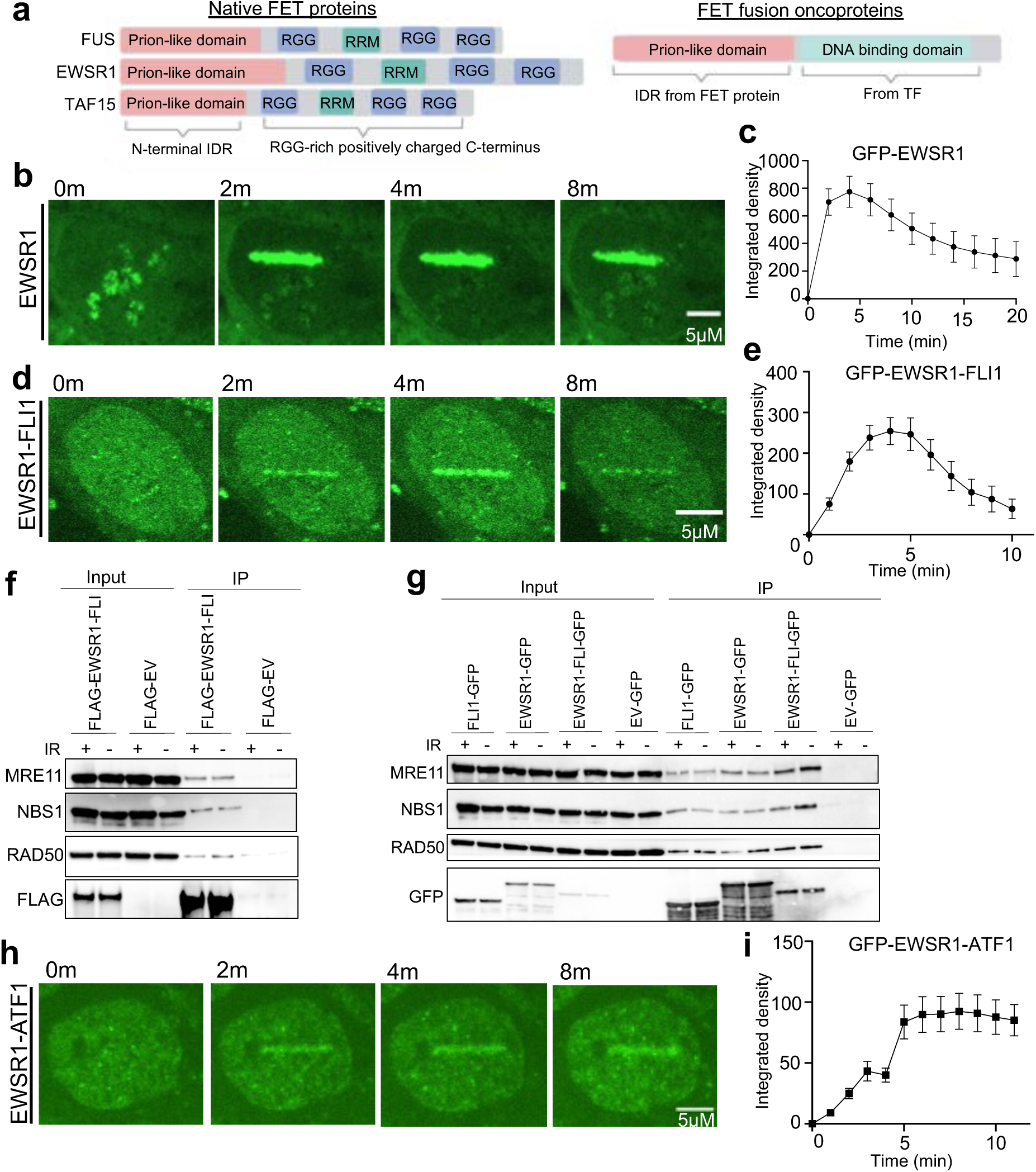
EWSR1 fusion oncoproteins are recruited to DNA DSBs. A) Structure schematic of FET proteins with N-terminal intrinsically disordered regions (IDR) and C-terminal Arginine-glycine-glycine (RGG) repeats, not present in FET fusion oncoproteins like EWSR1-FLI1. B-E, H, I) Representative images and quantification of protein accumulation at laser-induced DSBs in U2OS cells expressing either GFP-EWSR1 (B, C), GFP-EWSR1-FLI1 (D, E), or GFP-EWSR1-ATF1 (H, I). Quantification of at least 30 cells in 3 replicates. F, G) IP of FLAG and GFP tagged constructs (or EV) expressed in 293T cells +/- 5Gy IR, followed by Western blotting for MRN complex members. For all panels, error bars represent ± SEM, at least 3 replicates for each panel.

Importantly, EWSR1-FLI1 recruitment to DSBs represents an unanticipated neomorphic property of the fusion protein contributed by the N-terminal intrinsically disordered region (IDR) of EWSR1. Control experiments with the full-length FLI1 protein showed no accumulation at laser-induced DSBs (Fig. S5G). Consistent with the N-terminus of EWSR1 mediating DSB recruitment, mutation of the DNA binding domain of FLI1 (R2L2 mutant of EWSR1-FLI1^63^) had no effect on the DSB localization of EWSR1-FLI1 (Fig. S5H). Lastly, knockdown of native EWSR1 (using siRNA’s targeting C-terminal sequences not present in the fusion) reduced accumulation of EWSR1-FLI1 at laser-induced DSB stripes, indicating that the localization of EWSR1-FLI1 to DSBs depends, at least in part, on native EWSR1 (Fig. S5I).

Given the aberrant localization of EWSR1-FLI1 to DNA DSBs, we asked whether EWSR1-FLI1’s effects on ATM might result from direct protein-protein interactions with critical DDR factors that promote ATM activation. We utilized a candidate-based approach and performed co-IP experiments using EWSR1-FLI1 as the bait, both at baseline and after IR treatment. Interestingly, we found that EWSR1-FLI1 demonstrates an IR-independent interaction with all 3 members of the critical ATM activating MRN complex (MRE11, RAD50, NBS1) using both GFP and FLAG tagged versions of EWSR1-FLI1 (Figs. 5F, G). We also compared EWSR1-FLI1 with full-length native EWSR1 and FLI1 to assess domain contributions to MRN complex binding. EWSR1-FLI1 demonstrated greater binding to MRN complex members than either native full-length EWSR1 or full-length FLI1 (note the higher relative expression/IP levels for EWSR1 and FLI1, and that differences are most prominent for MRE11, Fig. 5G). These data suggest that suppression of ATM signaling in ES may result from re-localization of EWSR1-FLI1 to DNA DSBs and interaction/interference with the MRN complex, though further studies are needed to define the exact nature (i.e., direct or indirect) and functional impact of EWSR1-FLI1’s binding to the MRN complex, as well as contributions from other EWSR1-FLI1 interactions (e.g., with native FET proteins) not tested here.

Finally, we tested the generalizability of the concept that N-terminal IDRs, as a shared structural feature of FET fusion oncoproteins (Fig. 5A), could promote aberrant DSB recruitment in other tumors within this class. EWSR1-ATF1 is the sole oncogenic driver of clear cell sarcoma (CCS) and contains the identical N-terminal IDR sequence as EWSR1-FLI1. The EWSR1-ATF1 fusion oncoprotein localized to laser-induced DSBs with delayed recruitment kinetics similar to EWSR1-FLI1, though with differences in departure timing (Figs. 5H, I). Like EWSR1-FLI1, reintroduction of either 1 or all 3 RGG-rich domains into EWSR1-ATF1 resulted in earlier DSB recruitment kinetics and higher levels of overall recruitment in an RGG dose-dependent manner (Fig. S5J).

Analogous to our findings in ES, CCS cells are dependent on HR factors PALB2 and BRCA1 for survival (Fig. S5K, L). Moreover, transient knockdown of EWSR1-ATF1 in a patient-derived CCS cell line increased IR-dependent downstream ATM signaling (phosphorylation of ATM targets CHK2 and KAP1), with either no effect (DNA-PK) or slight decrease (ATR) in activation of the other upstream DDR kinases (Figs. S5M-P). These data also reveal differences between FET fusion oncoproteins which may relate to the efficiency of EWSR1-ATF1 knockdown in these experiments or to fusion-specific biology; only EWSR1-FLI1 affected the initial activation of ATM (autophosphorylation), while both EWSR1-ATF1 and EWSR1-FLI1 caused suppression of downstream ATM signaling. Despite these differences, the consequence of partial suppression of ATM signaling was a similarly increased reliance on the compensatory ATR signaling axis, as CCS cells displayed EWSR1-ATF1-dependent, nanomolar range sensitivity (IC50: 27nM) to ATR inhibitor treatment with elimusertib (Fig. 5SQ). Taken together, our results support a model in which EWSR1 fusion oncoproteins are aberrantly recruited to DSB repair sites and impair ATM function.

### Anti-tumor activity of ATR inhibitor elimusertib across FET rearranged PDX models

Lastly, we tested ATR inhibition as a therapeutic strategy in FET fusion oncoprotein-driven cancers. We hypothesized that the ATM/ATR synthetic lethality seen in ATM mutant (loss of function) cancers^56–58^ might extend to FET rearranged tumors with partial ATM defects caused by the FET fusion oncoproteins themselves. First, we determined whether FET fusion oncoprotein-driven cancer cell lines display increased sensitivity to the ATR inhibitor elimusertib that is currently being tested in multiple early phase clinical trials^64^. Our panel included 4 ES cell lines with varying TP53 and STAG2 status (see Supplementary Note 3), 1 CCS (EWSR1-ATF1), 2 desmoplastic small round cell tumor (EWSR1-WT1), and 1 myxoid liposarcoma cell line (FUS-CHOP), and multiple non-FET rearranged cancer cell lines as controls. FET fusion-driven cancer cell lines were significantly more sensitive to ATR inhibitor treatment than the control cell lines tested, with IC50’s between 20-60 nM (Fig. 6A) comparable to previously reported elimusertib IC50’s for ATM mutant cancer cell lines^65^. To test our hypothesis in a larger unselected panel of cancer cell lines, we utilized DepMap screening data for elimusertib in 880 cancer cell lines which included 17 ES samples. ES cell lines were again significantly more sensitive to ATR inhibition than the larger cohort of cancer cell lines (Fig. 6B), though our data does not exclude alternate mechanisms of sensitivity to ATR inhibition in non-ES cells such as ATM and other DDR gene mutations as previously reported^58^.

**Figure 6:**
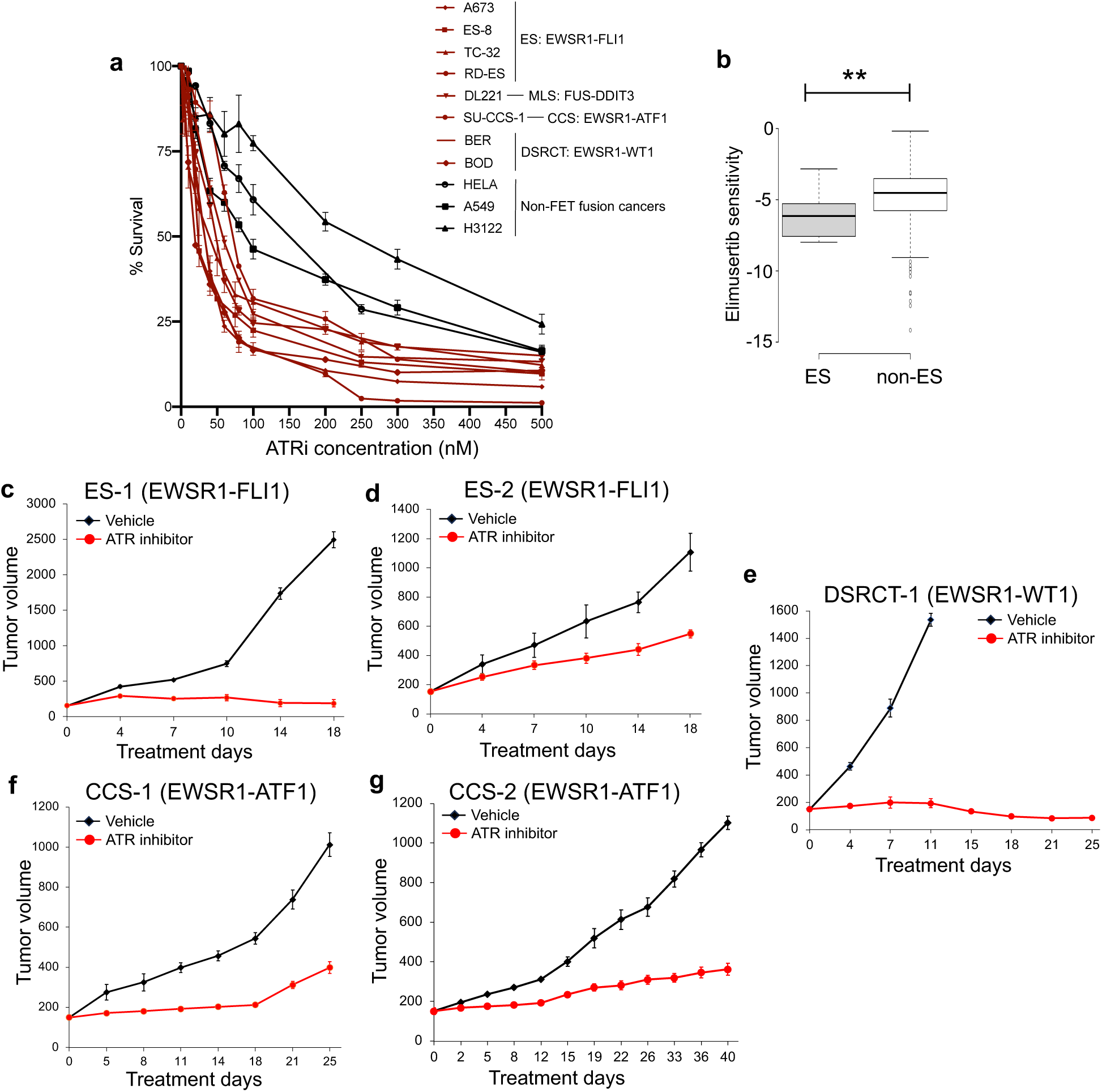
Anti-tumor activity of ATR inhibitor elimusertib in FET fusion PDXs. A) ATR inhibitor (elimusertib) dose-response curves for FET rearranged (red) and non-FET fusion driven cancer cell lines. MLS denotes myxoid liposarcoma. B) Analysis of DepMap Elimusertib sensitivity scores with samples divided into Ewing sarcoma (ES) and non-ES samples. ** denotes p<0.01 using unpaired t-test. C-G) Tumor volume measurements (mm^3^) of 5 FET rearranged PDX tumors treated with vehicle or elimusertib 40 mg/kg twice daily per oral gavage, on a 3 days-on/4 days-off schedule. All five models show significant anti-tumor responses with p<0.01 by unpaired T-test. For all panels, error bars represent ± SEM, at least 3 replicates in panel A, 4-8 mice per arm of PDX experiment.

We then tested whether the observed in vitro sensitivity to ATR inhibition would translate to in vivo patient-derived xenograft (PDX) models of FET rearranged cancers. Elimusertib was tested in 5 PDX models: 2 ES models (ES-1 from initial diagnosis and ES-2 from a multiply relapsed patient), 2 CCS models (CCS-1 from initial diagnosis and CCS-2 from relapsed disease), and 1 desmoplastic small round cell tumor (DSRCT) PDX derived from a post-treatment surgical resection, for which we also verified that the DSRCT fusion EWSR1-WT1 is recruited to laser-induced DSBs with similar kinetics to EWSR1-FLI and EWSR1-ATF1 (Figs. S6A, B). Clinical details on the tumor models are provided in Supplementary Table 2. Elimusertib was dosed at 40 mg/kg twice daily per oral gavage, on a 3 days-on/4 days-off schedule until tumors exceeded prespecified size endpoints as previously described^65^. There was minimal toxicity associated with this treatment regimen as assessed by weights and general activity scores, suggesting good tolerability of this dosing schedule.

Single-agent ATR inhibitor treatment significantly decreased tumor growth and increased progression-free survival in all 5 PDX models including those derived from relapsed ES and CCS patients (Figs. 6C-G). The DSRCT xenograft showed the best response (partial by RECIST criteria, >50% reduction in tumor volume, Fig. 6E). Interestingly, the ES-1 and CCS-1 PDX models from initial diagnosis showed prolonged stable disease (18 days of stable disease compared to immediate progression in vehicle treated tumors at day 4) and better overall response to elimusertib than the ES-2 and CCS-2 PDXs derived from relapsed disease, though testing of additional ES and CCS PDX models will be important to clarify whether PDXs derived from diagnostic and relapsed disease have differential responses to ATR inhibition.

To identify pharmacodynamic markers of elimusertib activity, we performed immunohistochemistry (IHC) for Ki-67 (cell proliferation), cleaved caspase-3 (apoptosis), and DNA damage (gamma-H2AX, gH2AX). We observed decreased Ki-67 in elimusertib treated tumors compared to vehicle controls reflecting decreased proliferation, while increased cleaved-caspase 3 was only observed in the 2 ES PDXs (Figs. S6C-F). Consistent with prior reports^66,67^, we were unable to validate a direct biomarker of ATR inhibition as p-ATR could not reliably be assessed by IHC and its downstream target p-CHK1 did not show a significant difference between vehicle and elimusertib treatment (Figs. S6C-F), which may reflect the low-level of ATR pathway activation at baseline. However, gH2AX proved to be a reliable biomarker for elimusertib activity. The fraction of gH2AX positive cells was significantly increased in all elimusertib treated tumors compared to vehicle controls, consistent with unrepaired and ongoing DNA damage as the mechanism of cell death and anti-tumor activity. Taken together, our data demonstrate that FET rearranged cancers are preferentially sensitive to ATR inhibition and that single agent treatment with the ATR inhibitor elimusertib has in vivo anti-tumor activity across multiple FET fusion oncoprotein-driven PDX models.

## Discussion

Here we describe how an oncogene can disrupt physiologic DNA damage repair by suppressing ATM activation and downstream signaling. We establish the compensatory DDR kinase, ATR, as a collateral dependency and therapeutic opportunity to target the (partial) ATM defects induced by the oncoproteins themselves using multiple FET fusion-driven cell line and PDX models. These data provide a template for the broader utilization of DDR-directed therapies in cancer through improved understanding of oncogene-DDR network interactions.

The specific nature of the DNA repair defect in ES has been the subject of much debate given the strong clinical and laboratory-based data demonstrating chemo- and radio-sensitivity^11–15^. Our discovery that ES tumors are not “BRCA-like”, but instead harbor partial ATM defects, may help explain the lack of clinical responses to PARP inhibitor monotherapy in ES patients^16,68^ (unlike HR deficient BRCA mutant and BRCA-like cancers^17,18^). ES cells are dependent on key HR genes for survival and direct analysis of ES patient tumors revealed none of the genomic hallmarks of HR loss. Instead, we show that EWSR1-FLI1 suppresses ATM activation and downstream signaling in response to DNA damage (IR), without impacting the other apical DDR kinases (DNA-PK and ATR). While we do not exclude the possibility that additional DNA repair defects contribute to the DSB-sensitivity of ES cells, our finding of ATM (and not HR) impairment has important clinical implications for ongoing attempts to target the DDR in this disease, especially relating to ongoing PARP inhibitor combination trials^69^.

To contextualize the magnitude and biological relevance of the partial ATM defects reported here, we benchmarked EWSR1-FLI1 against loss of canonical ATM activators. The effect size of EWSR1-FLI1 mediated suppression of ATM activation/signaling is comparable to loss of established ATM activators including MDC1^49^, RNF8^51^ or RNF168^54^ (whose loss confer significant radio-sensitivity to IR), but only ∼30-60% of the near-complete functional ATM deficiency seen with loss of the MRN complex. Thus, the magnitude of ATM impairment in ES is biologically significant, but not absolute. We further demonstrate the functional importance and therapeutic relevance of partial ATM suppression through increased reliance on a compensatory DDR kinase, ATR, and EWSR1-FLI1-dependent synthetic lethality with inhibitors of ATR and its major downstream target CHK1. The observed ATM effects do not appear to depend on TP53 or STAG2 status as the phenotypes were observed in both mutant and wild-type ES cell lines (see Supplemental Note 3 for additional discussion). We note that the genomic instability of ATM mutant (loss of function) tumors has not been associated with a specific mutational or copy number alteration (CNA) signature^70^ thus precluding comparison here, though ES tumors do display recurrent CNAs (trisomy 8 in ∼50% of cases, ∼20% with trisomy 12 or 1q gain^34^) which warrant further study. In total, our data nominate partial suppression of ATM as a therapeutically targetable DNA repair lesion in ES.

How does EWSR1-FLI1, a TF fusion oncoprotein, create these DNA repair defects in ES? We did not identify a set of EWSR1-FLI1 transcriptional targets that explain these ATM effects (e.g., MRN complex genes were not transcriptionally suppressed nor absent at the protein level by Western blotting). Instead, we find that EWSR1-FLI1 is recruited to DNA DSBs through homotypic intrinsically disordered region (IDR) domain interactions with native EWSR1. Moreover, we show that EWSR1-FLI1 directly interacts with all three members of the MRN complex that is essential for ATM activation. Both DSB recruitment and MRN complex binding represent neomorphic functions of the fusion protein contributed by the EWSR1 N-terminal IDR, thus distinguishing EWSR1-FLI1 from wild-type, full-length FLI1. The specific nature of EWSR1-FLI1’s interaction with the MRN complex (e.g., direct or indirect, effect on MRN complex function) and whether other DNA repair proteins interact with EWSR1-FLI1 and contribute to the observed ATM effects (e.g., native EWSR1) are important unanswered questions for future studies to address. In summary, we show that EWSR1-FLI1 is unexpectedly recruited to DNA DSBs and interacts with the MRN complex, which may provide a molecular basis for the partial ATM defects in ES that we describe here.

More generally, the shared structural organization across the class of FET fusion oncoproteins (i.e., retention of N-terminal IDRs that mediate homotypic and heterotypic FET protein interactions and loss of C-terminal RGG domains that promote rapid DSB recruitment, Fig. 5A) raised the intriguing hypothesis that FET fusion-driven cancers harbor a common set of DNA repair defects. Indeed, we show that the clear cell sarcoma fusion oncoprotein EWSR1-ATF1 localizes to DNA DSBs and induces similar suppression of IR-dependent ATM signaling (though with differences in ATM autophosphorylation that may relate to efficiency of EWSR1-ATF1 knockdown, Fig. S5M). Analogous to our findings in ES, CCS cells also display FET fusion oncogene-dependent nanomolar range sensitivity to the ATR inhibitor elimusertib. These data prompted us to test ATR inhibition across the class of FET rearranged cancers as a synthetic lethal therapeutic strategy to target partial ATM defects induced by FET fusion oncoproteins.

We selected a potent ATR inhibitor in clinical development (elimusertib) and observed nanomolar range IC50’s and preferential sensitivity of FET fusion-driven cancer cell lines in vitro (IC50 range: 20-60nM, comparable to reported elimusertib IC50’s in cell lines with pathogenic loss of function ATM mutations^65^). Moreover, we tested single agent elimusertib treatment in 5 PDX models including 2 ES, 2 CCS, and 1 DSRCT model. All 5 models showed significant decreases in tumor growth and prolongation of progression-free survival including ES and CCS PDXs from multiply relapsed patients. Given that PDXs are a stringent bar in terms of disease control due to their inherent heterogeneity and single agents are highly unlikely to be curative for these set of diseases, the ability of ATR inhibitor monotherapy to control tumor growth in all 5 models was a striking finding. Our findings are consistent with published reports testing elimusertib in pediatric cancer PDX’s (ES was the only FET rearranged tumor tested)^71^, but differ from a recent report with another ATR inhibitor in clinical trials (berzosertib) that showed no anti-tumor activity as monotherapy in ES cell line xenografts^72^, suggesting important potential differences between ATR inhibitors that warrant future study. We note that previously described mechanisms of ATR inhibitor sensitivity such as PGBD5 transposase expression^73^ or replication stress^15^ could also contribute to the elimusertib effects that we observed. In total, these data provide strong preclinical rationale for specifically testing elimusertib (and combinations thereof) in FET rearranged cancers as part of ongoing early phase clinical trials.

Finally, what might be the selective advantage for an oncogene to also suppress physiologic DNA repair? We speculate here that by partially disabling DDR signaling, cells with FET fusion oncoproteins can tolerate high levels of DDR activation caused by transcription and replication stress induced by the oncoprotein itself and thus overcome an important barrier to cellular transformation. Whether other TF fusion oncoproteins (or oncogenes more generally) interact with or disrupt tumor suppressive DDR signaling networks as part of oncogenesis will be an intriguing topic for future work. In summary, our study describes how growth-promoting oncogenes can also interfere with physiologic DNA damage repair and provides rationale for a new targeted therapeutic strategy in ES and the broader class of undruggable FET fusion-driven cancers.

---

**Supplemental Figure 1:**
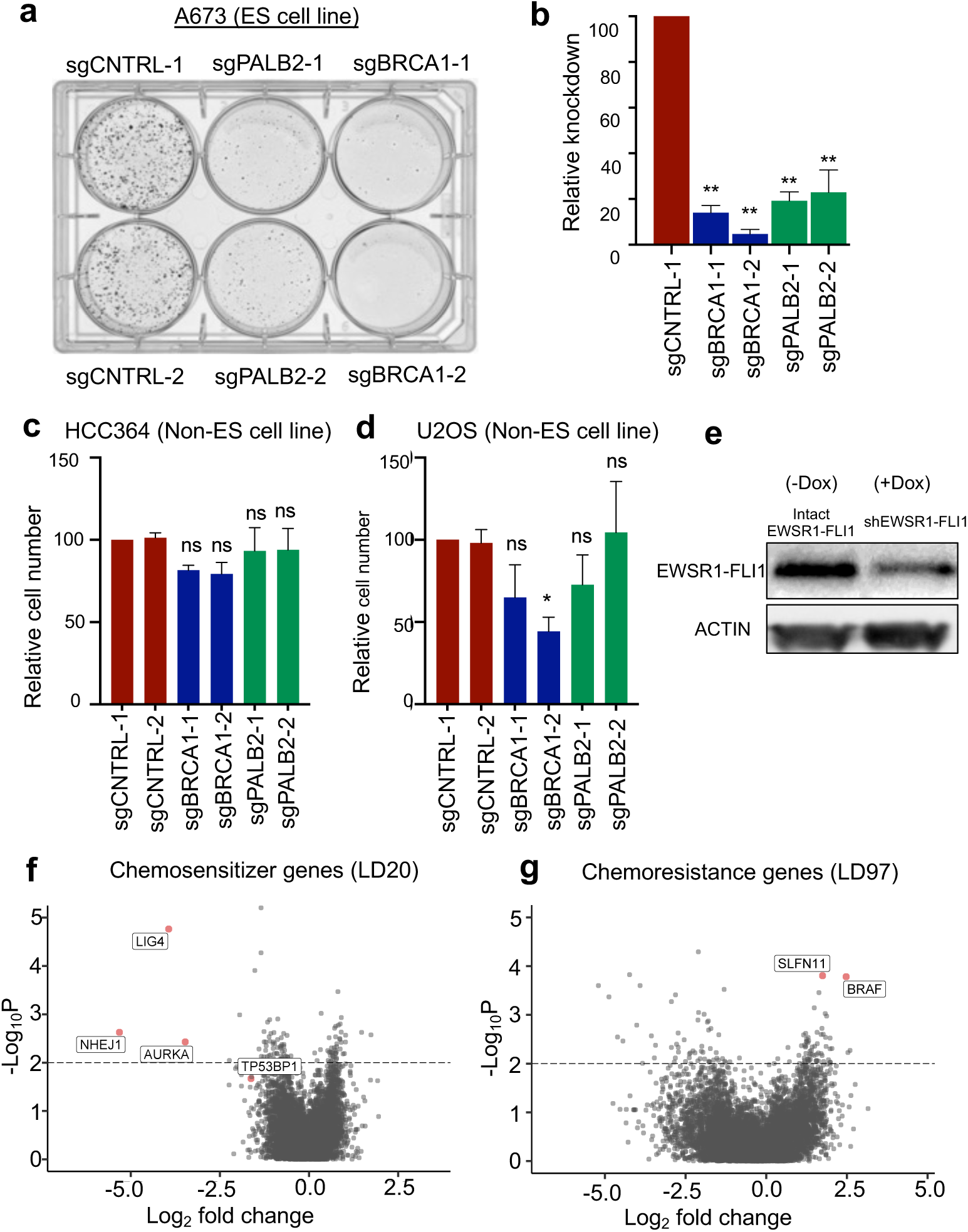
ES cells are dependent on specific HR factors for survival. A) Representative crystal violet staining of 72-hour growth assay in dCas9 expressing A673 ES cells upon introduction of 2 independent sgRNAs against BRCA1, PALB2, or control. B) Relative knockdown of each sgRNA by qPCR, n = 3. C, D) Growth assays in dCas9 expressing cancer cell lines upon introduction of 2 independent sgRNAs against BRCA1, PALB2, or control, n = 4, 4. E) Western blot of EWSR1-FLI1 levels in A673 with doxycycline-inducible shRNA against EWSR1-FLI1. F, G) Volcano plot of sgRNA log_2_fold change relative to DMSO control (X-axis) vs -log10(p value). Full screen results provided in Supplementary Table 1. For all panels, * denotes p< 0.05, ** denotes p < 0.01, ns denote not significant by one-way ANOVA with post hoc Tukey’s HSD test.

**Supplemental Figure 2:**
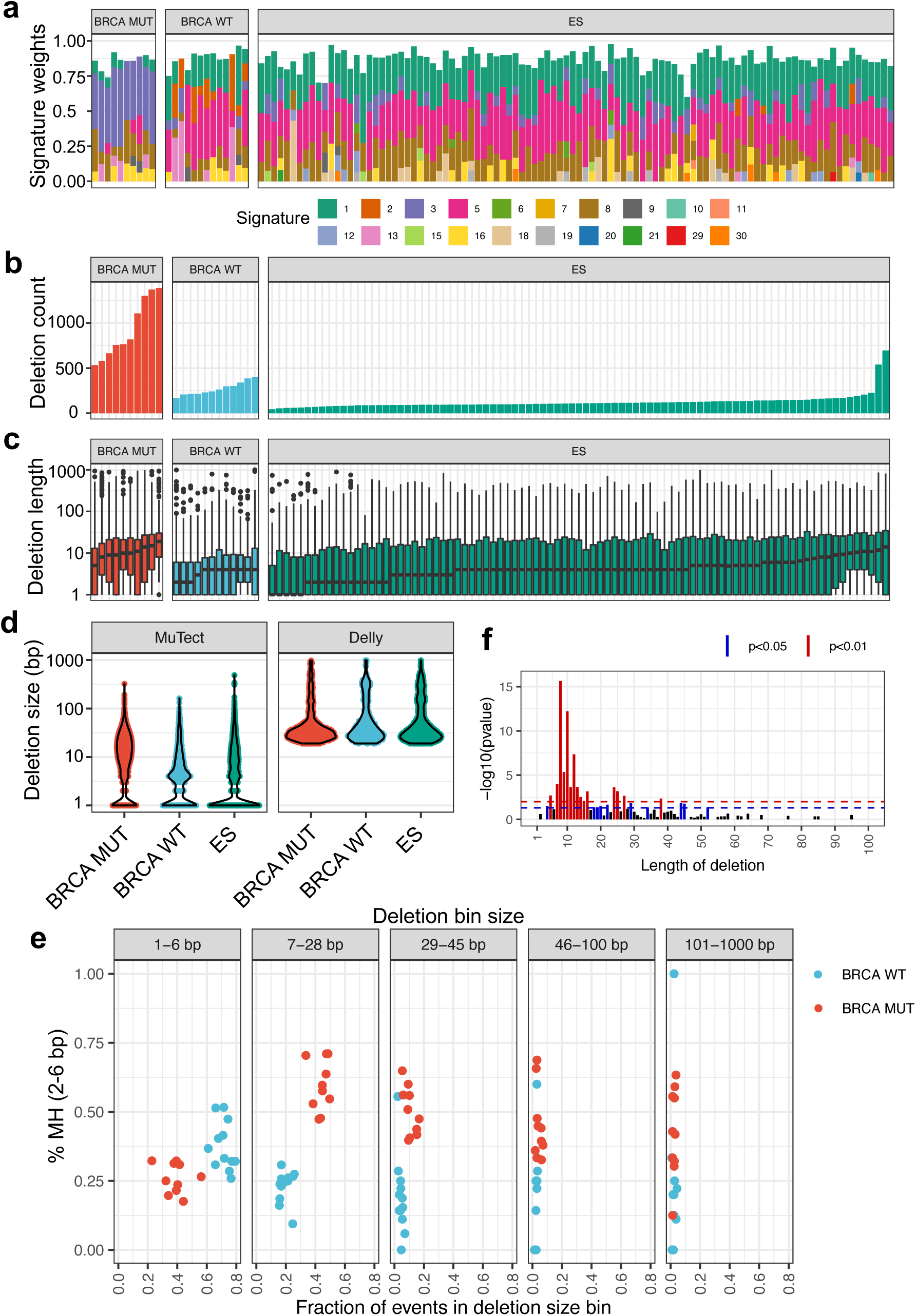
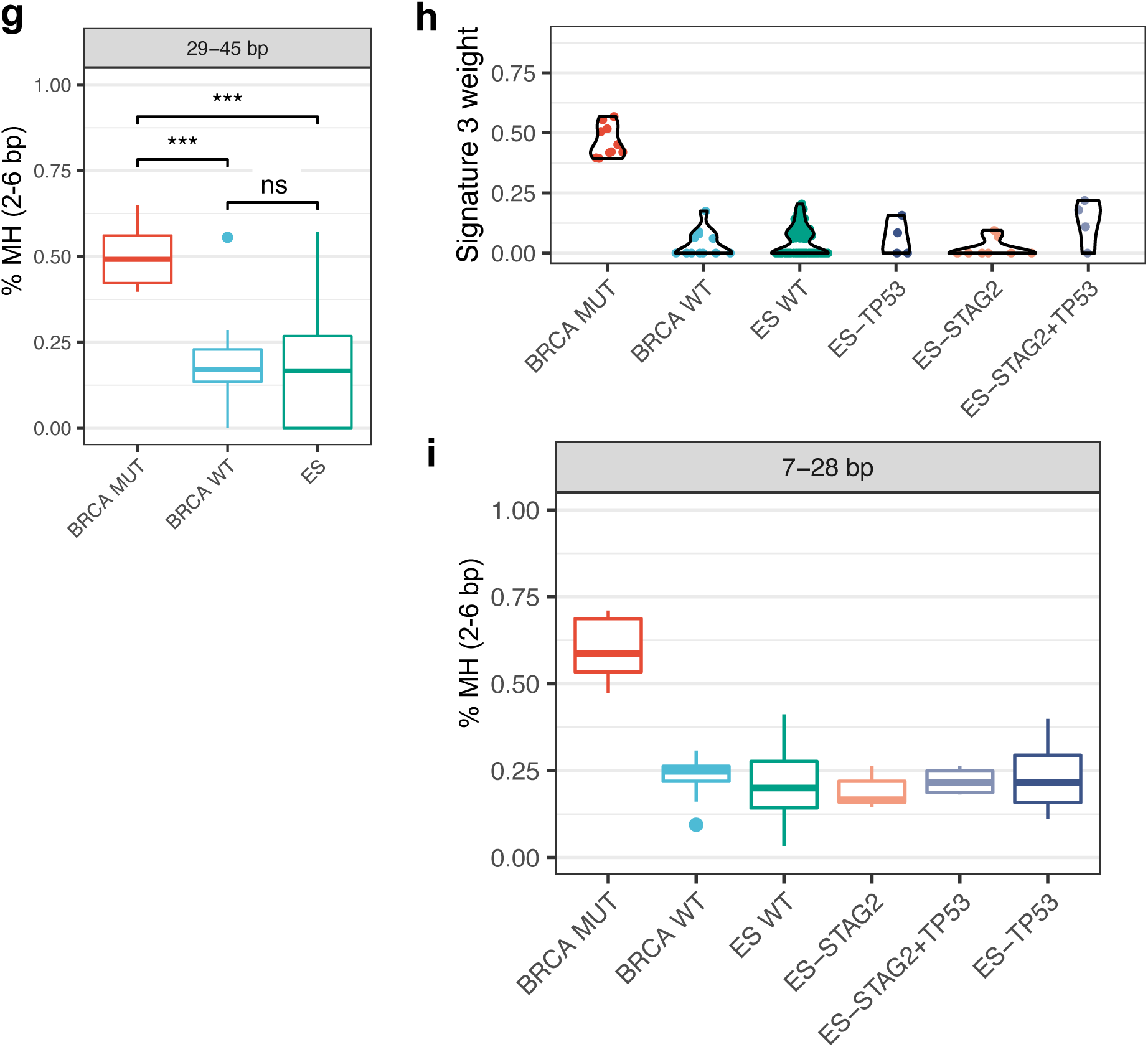
ES patient tumors do not display the genomic scars of HR deficiency. A) Mutational signature breakdown for BRCA mutant (MUT)/wild-type (WT) and ES tumors. B) Total number of unique deletions per sample. C) Deletion length per sample. D) Deletion size per tumor sample shown individually for MuTect and Delly deletion calls. E) Deletion size/microhomology (MH) profile for only BRCA WT and BRCA MUT samples showing separation between the groups present in the 7-28bp and 29-45bp bins in both the fraction of events in deletion size bin (X-axis) and percentage of MH (Y-axis). F) Graph of Fisher test significance (p value) for deletion lengths and MH status. G) Breakout of the 29-45 base pair (bp) deletion length bin showing the fraction of deletions with 2-6 bp microhomology. H, I) Signature 3 weight (H) and 2-6bp breakpoint MH in the 7-28bp deletion bin (I) in ES samples subdivided by TP53 and STAG2 mutational status. ES WT are wild-type for p53 and STAG2. For all panels, *** denotes p< 0.001 by Mann-Whitney test, ns denotes not significant. European Genome-phenome Archive accession numbers: ES samples (EGAS00001000855 - Institute Curie cohort and EGAS00001000839 - St. Jude’s cohort)^34^, breast cancer samples (EGAD00001001322)^35^.

**Supplemental Figure 3:**
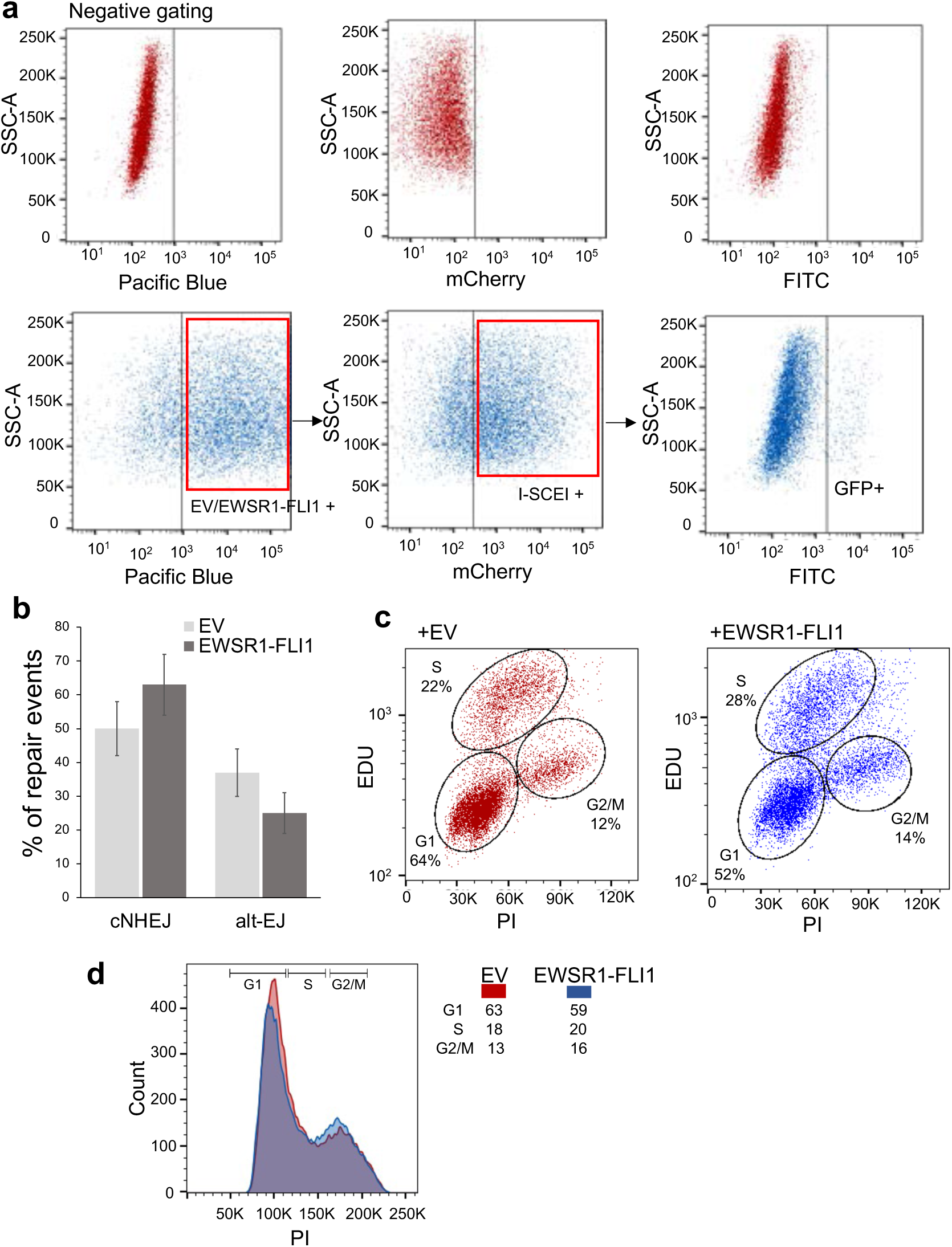
EWSR1-FLI1 reduces resection-dependent DSB repair. A) Gating schema for pathway-specific DSB repair reporter assays in U2OS cells using dual promoter vector expressing mTagBFP and EWSR1-FLI1 (or empty vector, EV), followed by introduction of mCherry-I-SceI endonuclease. GFP positivity is measured at 72 hours. B) Sequence analysis of repair events after transient introduction of I-SceI DSB reporter (NHEJ- I)^40^ in U2OS cells expressing EV or EWSR1-FLI1. Error bars represent ± SEM, p<0.05 for both comparisons, paired t-test, n=3. C) Cell cycle profiles using EdU incorporation in U2OS cells +/- EWSR1-FLI1. D) Cell cycle profiles using Propidium Iodide (PI) +/- EWSR1-FLI1.

**Supplemental Figure 4:**
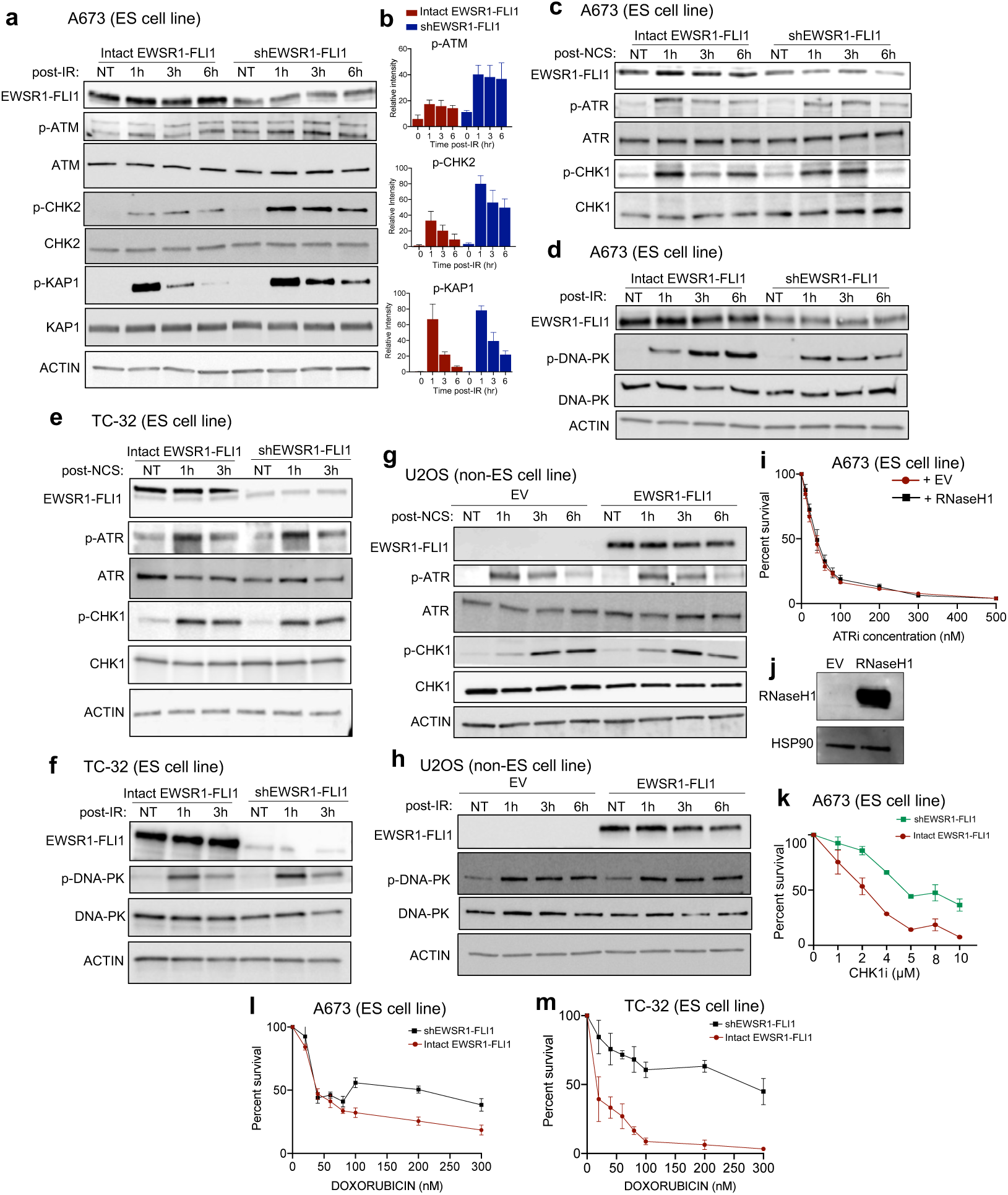
ATM activation and signaling is suppressed by EWSR1-FLI1. A, B) Western blotting and quantification upon 5 Gy IR at indicated time points in ES cell line A673 with doxycycline-inducible shRNA against EWSR1-FLI1. C-H) Western blotting upon 200 ng/ml NCS treatment (C, E, G) or 5 Gy IR (D, F, H) at indicated time points in A673 and TC-32 ES cells with dox-inducible shRNA against EWSR1-FLI1 or upon EV or EWSR1-FLI1 expression in non-ES (U2OS) cancer cell line. I, J) ATR inhibitor (elimusertib) dose-response curves for ES cell line A673 +/- RNAseH1 overexpression and confirmatory Western blot of RNAseH1 expression (J). K) CHK1 inhibitor (LY2603618) dose-response curves in A673 with dox-inducible shRNA against EWSR1-FLI1, p<0.05 by paired t-test, n=3. L, M) Doxorubicin dose-response curves for ES cell lines A673 and TC-32 with dox-inducible shRNA against EWSR1-FLI1, p<0.05 for both comparisons, paired t-test, n=3. For all panels, error bars represent ± SEM and represent at least 3 replicates for each panel.

**Supplementary Figure 5:**
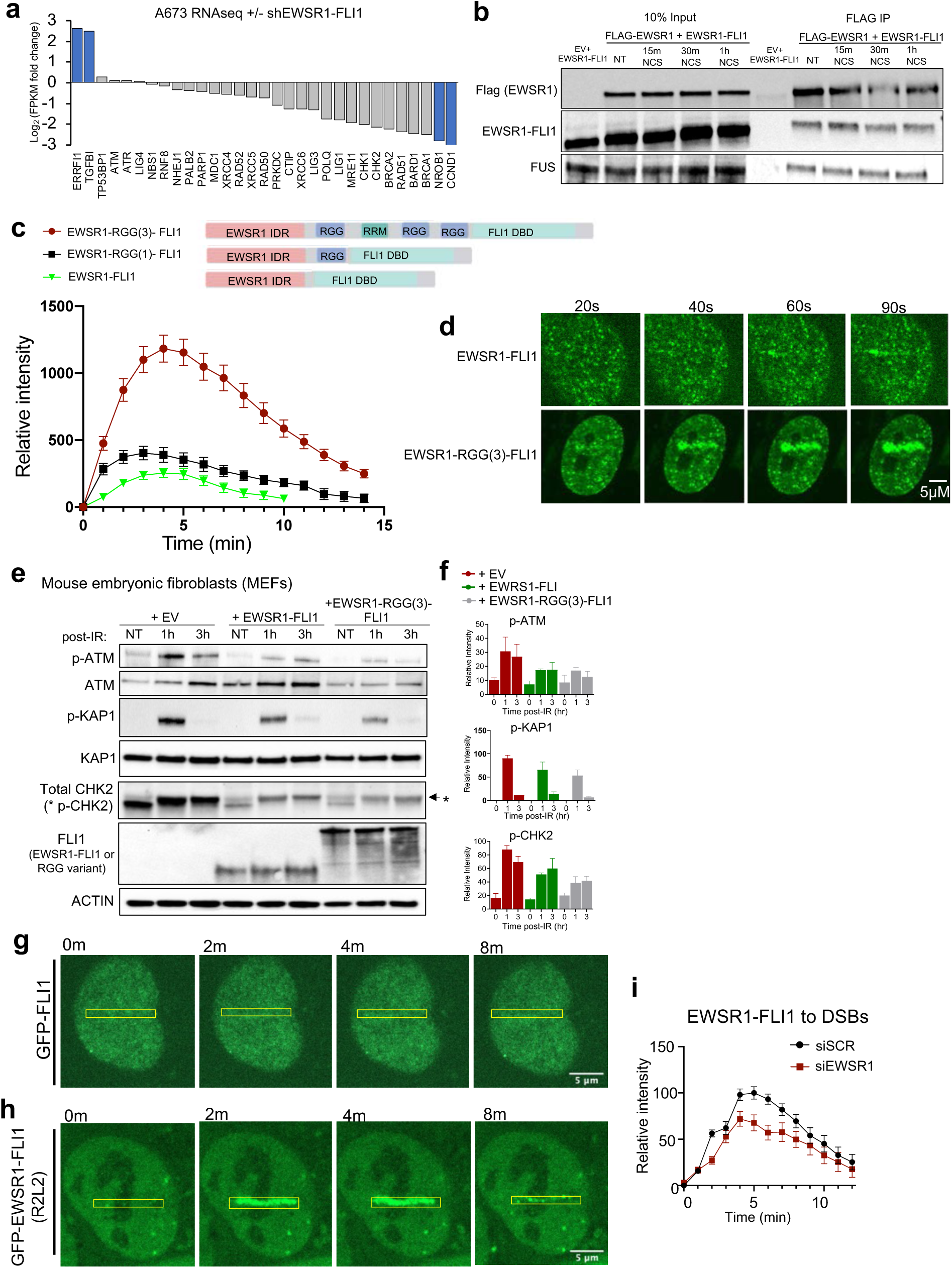
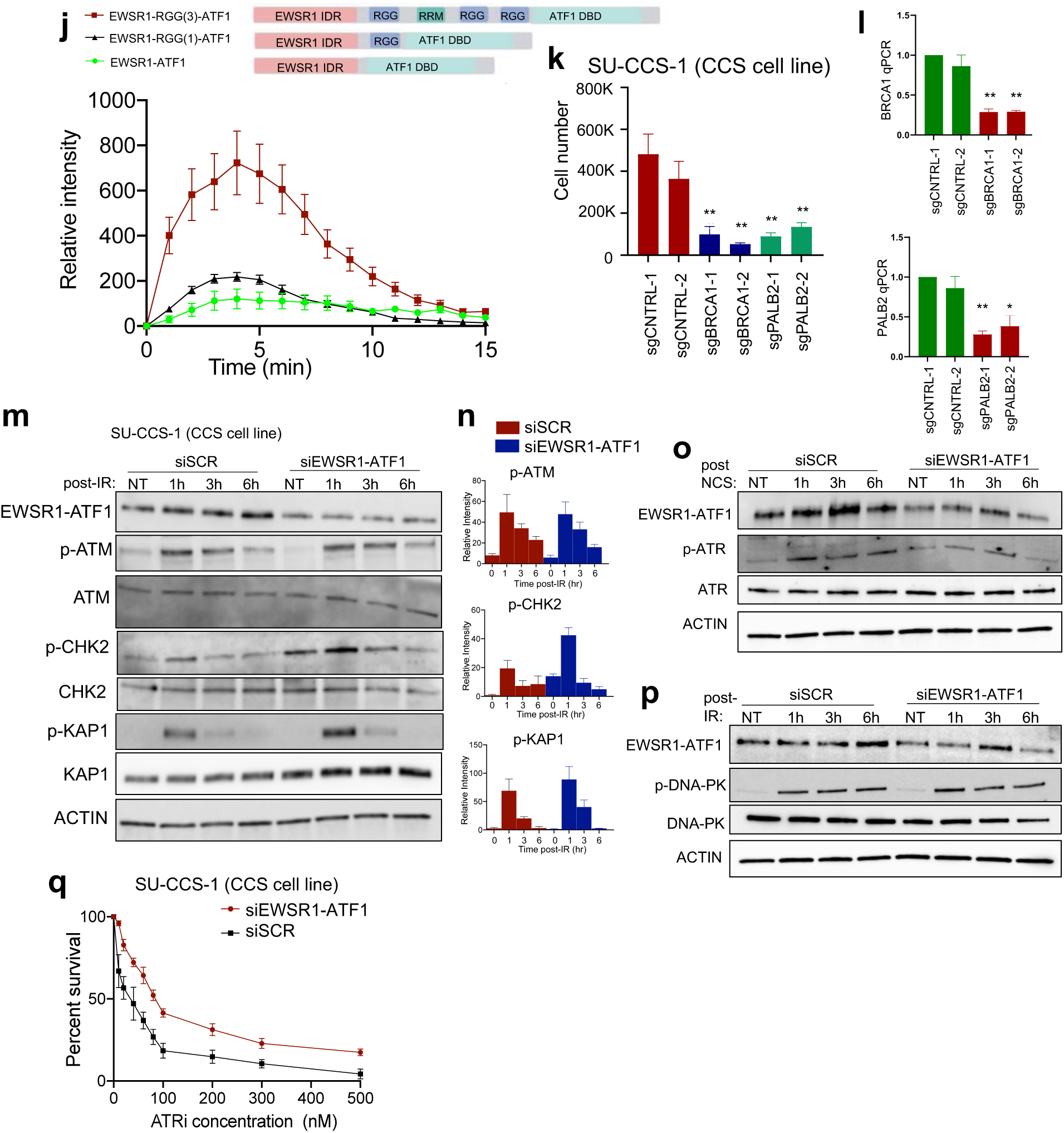
EWSR1 fusion oncoproteins are recruited to DNA DSBs. A) Analysis of published RNAseq data^60^ in A673 ES cells with EWSR1-FLI1 knockdown. Blue denotes canonical EWSR1-FLI1 targets. Negative values are genes upregulated by EWSR1-FLI1. B) Immunoprecipitation (IP) of FLAG-tagged EWSR1 expressed in 293T cells +/- EWSR1-FLI1. NCS treatment at 200 ng/ml. C, D) Structure schematic of RGG containing versions of EWSR1-FLI1. DBD denotes DNA-binding domain. Representative image and quantification of laser-induced DSB accumulation of GFP-tagged constructs in U2OS cells. E, F) Western blotting and quantification after IR (5 Gy) at indicated time points in MEFs infected with EV, EWSR1-FLI1, or EWSR1-RGG(3)-FLI1. G, H) Laser micro-irradiation of GFP-tagged full-length FLI1 or DNA binding deficient mutant (R2L2) of EWSR1-FLI1. I) Quantification of GFP-EWSR1-FLI1 accumulation at laser-induced DSBs in a non-ES cell line (U2OS) upon siRNA against native EWSR1 (targeting C-terminal sequences) or scramble (siSCR) for 72 hours. n= 28, 27 cells, p<0.01 using paired t-test. J) Structure schematic of RGG containing versions of EWSR1-ATF1 and quantification of laser-induced DSB accumulation of GFP-tagged constructs in U2OS cells. K) Growth assays in dCas9 expressing CCS cell line SU-CCS-1 upon introduction of 2 independent sgRNAs against BRCA1, PALB2, or control. n = 4. L) Relative gene knockdown of each sgRNA by qPCR in SU-CCS-1 cells, n = 3. M, N) Western blotting and quantification after IR (5 Gy) in CCS cell line SU-CCS-1 upon 72 hour siRNA treatment against EWSR1-ATF1 or Scramble (siSCR) control. O, P) Representative Western blots upon 200 ng/ml NCS treatment (O) or 5 Gy IR (P) in CCS cell line (SU-CCS-1) after 72-hour siRNA treatment against EWSR1-ATF1 or scramble (SCR) control. Q) ATR inhibitor (elimusertib) dose-response curves for CCS cell line SU-CCS-1 upon siRNA against EWSR1-ATF1 or siSCR, p<0.05 by paired t-test, n=3. For all panels, error bars represent ± SEM, at least 3 replicates for each panel. ** denotes p < 0.01

**Supplemental Figure 6:**
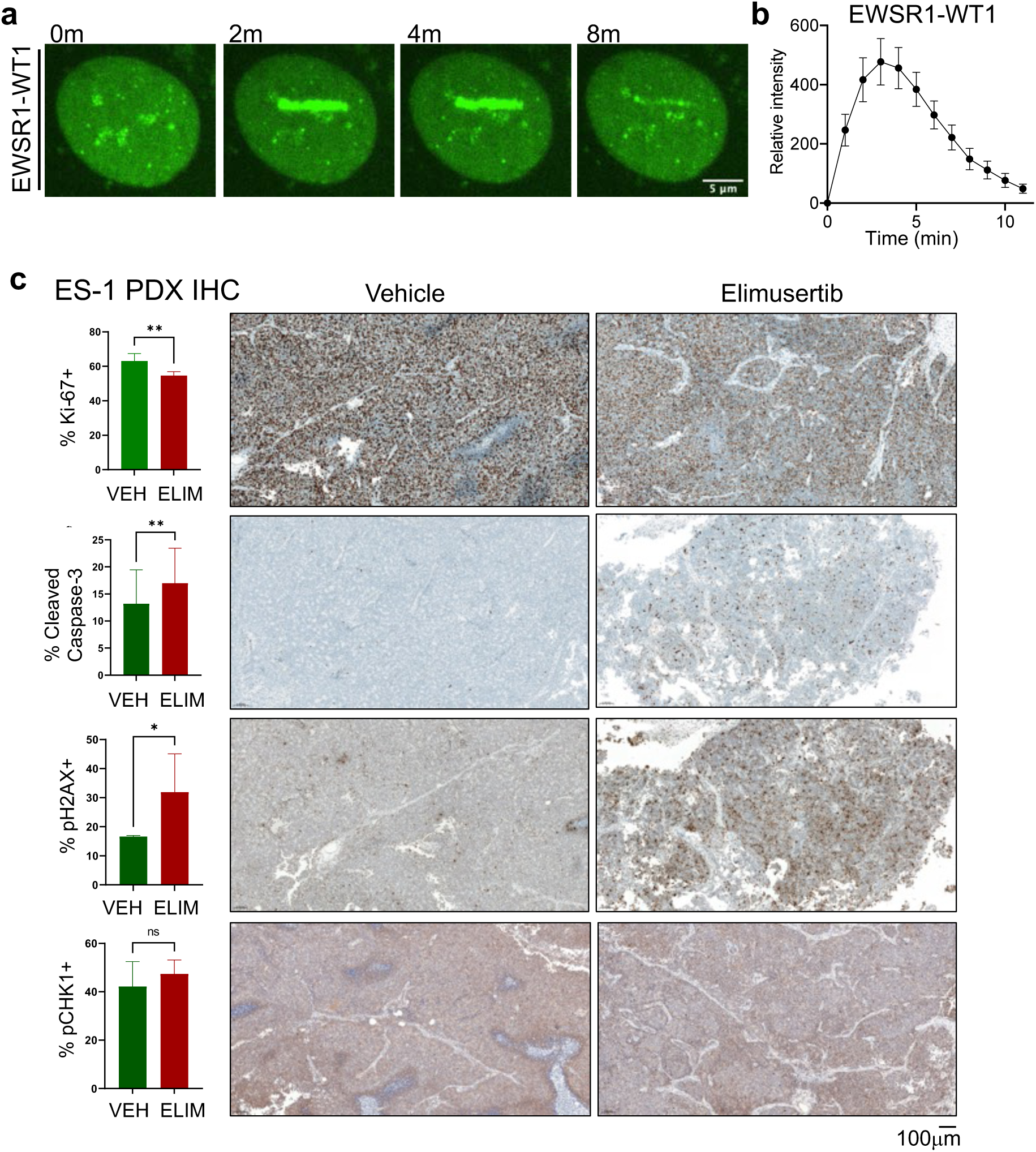
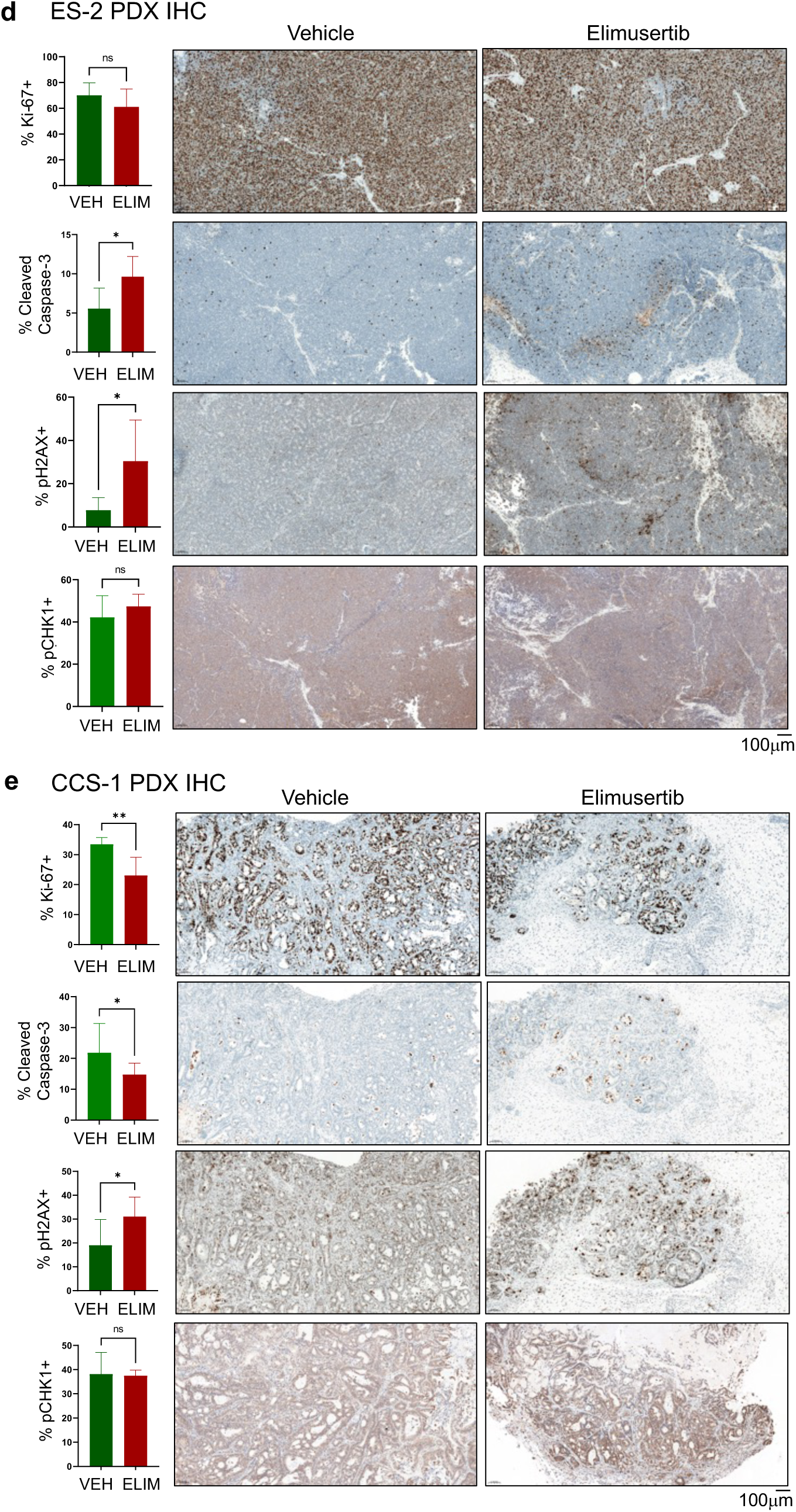
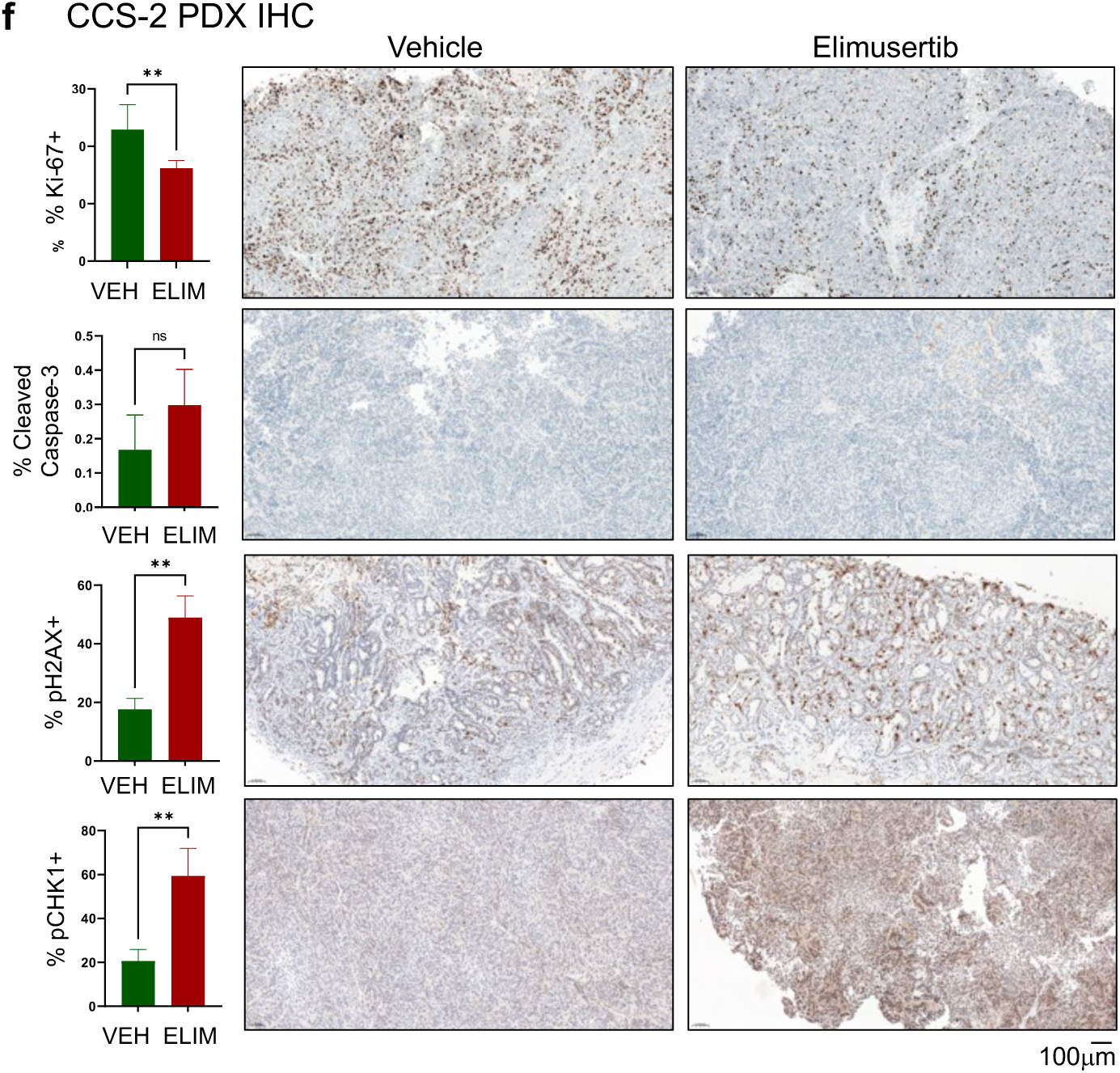
Anti-tumor activity of ATR inhibitor elimusertib in FET fusion PDXs. A, B) Representative images and quantification of protein accumulation at laser-induced DSBs in U2OS cells expressing GFP-EWSR1-WT1. Quantification of at least 30 cells in 3 replicates. C-F) Immunohistochemistry (IHC) staining quantification and representative fields of PDX tumors at end of therapy. VEH refers to vehicle treated tumors; ELIM are elimusertib treated. Automated quantification using QuPath. Note continued regression of DSRCT PDX precluded IHC analysis. Error bars represent ± SEM, * denotes p<0.05, ** denotes p<0.01, unpaired T-test.

**Supplemental Figure 7:**
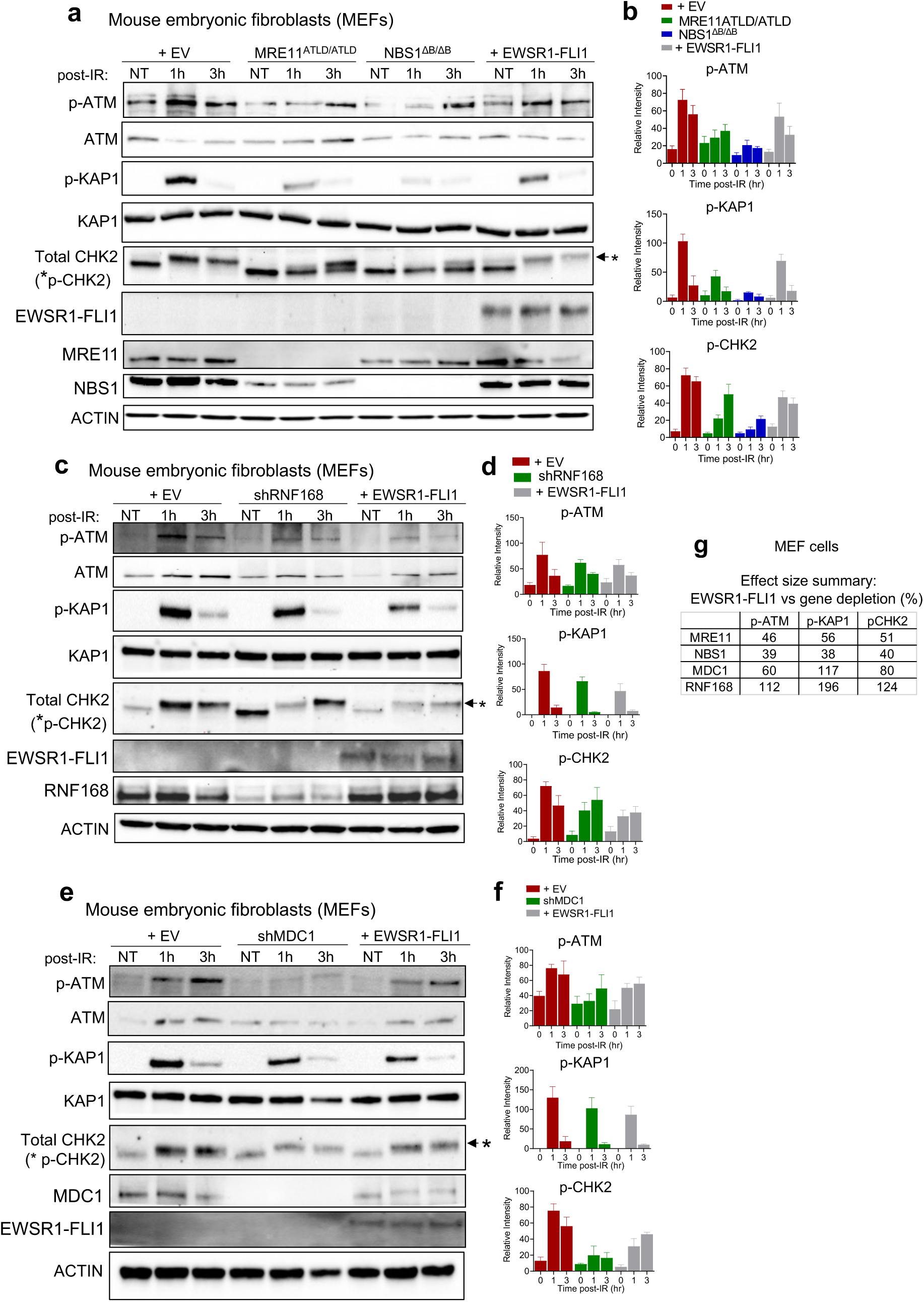
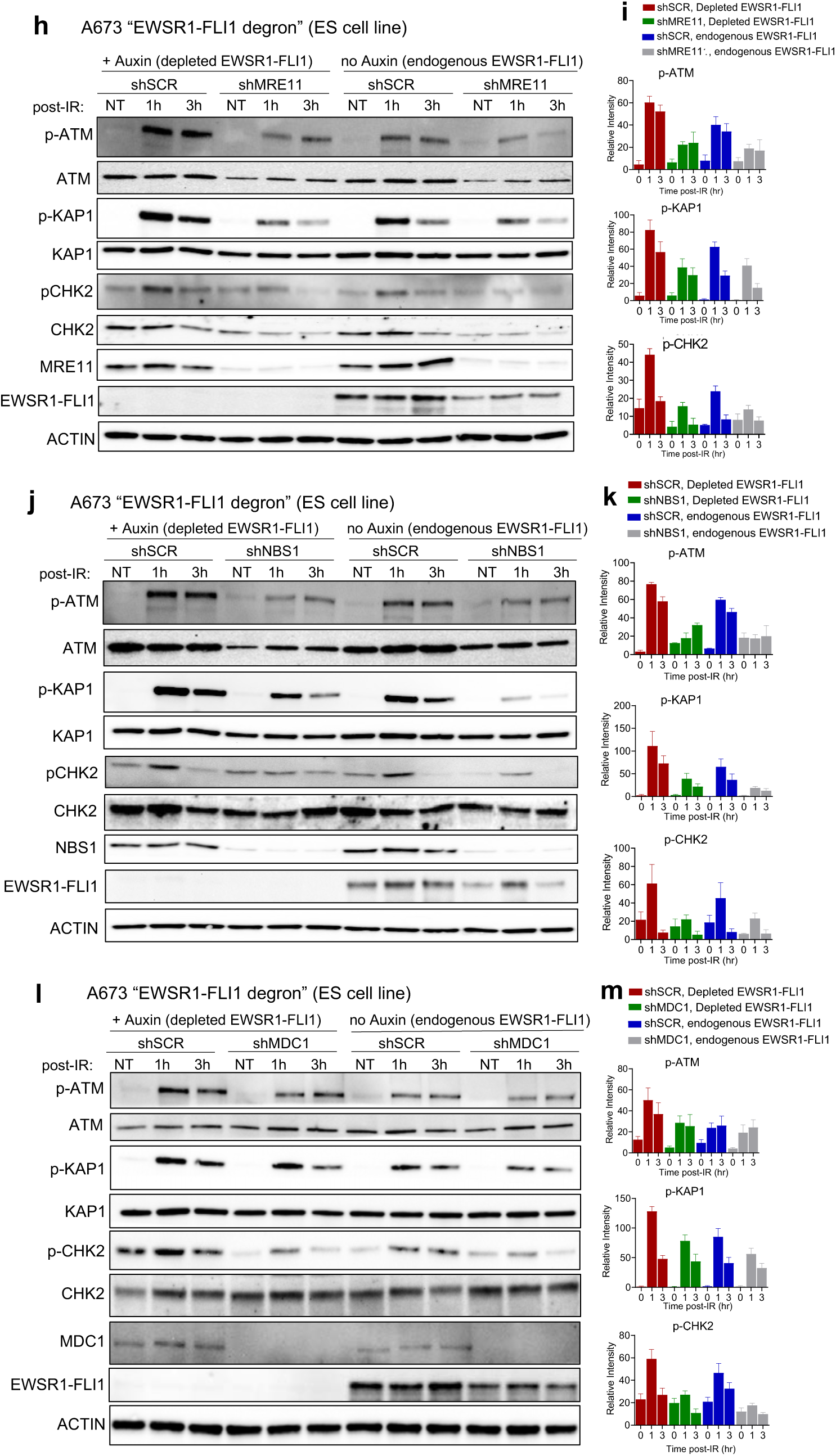
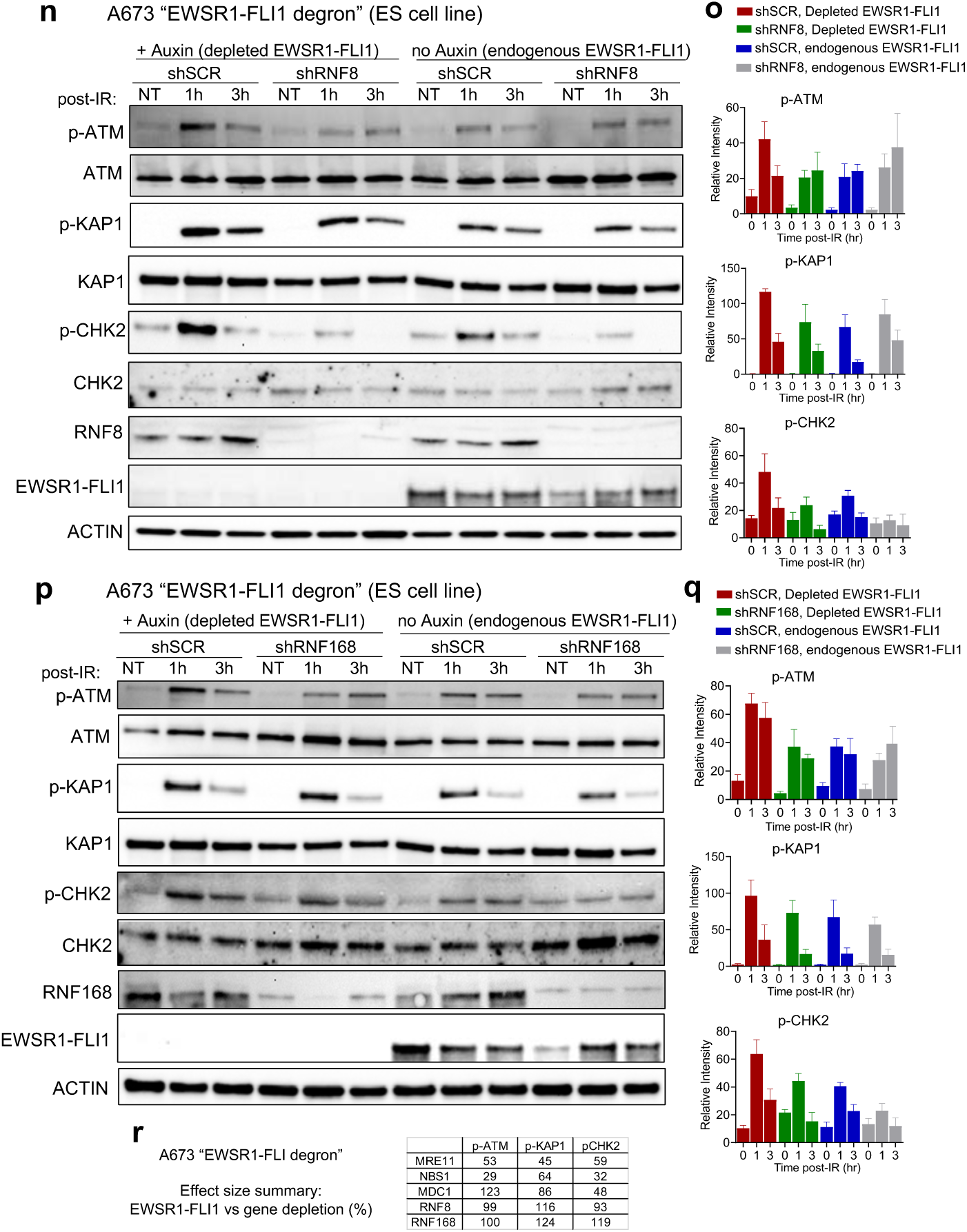
Comparison of EWSR1-FLI1 to loss of canonical ATM activators. A-F) Western blotting and quantification after IR (5 Gy) at indicated time points in either wild-type (WT) MEFs with empty vector (EV) or EWSR1-FLI1 expression, MRE11^ATLD/ATLD^ and NBS1^ΔB/ΔB^ MEFs (A, B), or WT MEFs with an shRNA against RNF168 or MDC1. G) Effect size comparison in MEFs of EWSR1-FLI1 to loss of ATM regulator in terms of ATM suppression (p-ATM/p-CHK2/p-KAP1) based on 1hr time point post-IR. % represents ratio of reduction from baseline (EV) of EWSR1-FLI1 vs. knockdown of ATM regulator. H-Q) Western blotting and quantification upon 5 Gy IR in ES cell line A673 “EWSR1-FLI1 degron” infected with shRNAs against MRE11, NBS1, MDC1, RNF8, RNF168, or shSCR. 200 μM IAA (auxin) added 3 hours prior to IR to degrade endogenous EWSR1-FLI1. R) Effect size comparison in A673 “EWSR1-FLI1 degron” cells of endogenous EWSR1-FLI1 to loss of ATM regulator in terms of ATM suppression based on 1hr time point post-IR. % represents a ratio of reduction from maximal ATM activity (gene knockdown, EWSR1-FLI1 depleted: red bars) for (1) shScramble (shSCR), endogenous EWSR1-FLI1 (blue bars) versus (2) gene knockdown, EWSR1-FLI1 depleted (green bars). For all panels, error bars represent ± SEM and represent at least 3 replicates for each panel.

## Supplementary Note 1

We compared our CRISPRi screening results with the DepMap project cancer cell line database (https://depmap.org/portal)^74^. Our analysis shows a statistically significant, but modest, difference in the distribution of PALB2 dependence scores when comparing Ewing sarcoma (ES) cell lines with the larger non-ES cancer cell line database, but no significant difference for BRCA1. We hypothesize that these differences may reflect the challenge of studying “essentiality” for double-strand break (DSB) repair genes using a catalytically active Cas9 nuclease that generates DSBs, which was our initial rationale for selecting a CRISPRi-based approach.

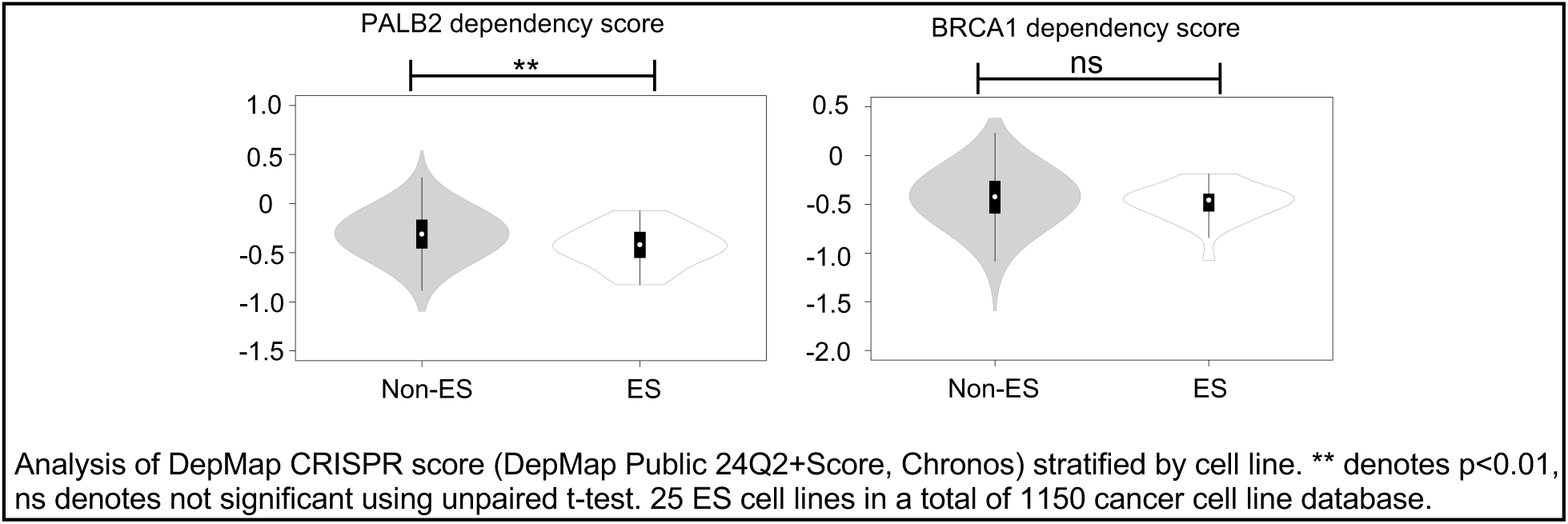

In support of this hypothesis, we noted a large discrepancy in the essentiality of these homologous recombination (HR) genes between CRISPR/Cas9 nuclease and RNAi screens in the DepMap database, with a much higher percentage of cancer cell lines found to be dependent on PALB2 and BRCA1 when assayed using a catalytically active Cas9 nuclease.

| Gene | % Dependent Cancer Cell Lines in CRISPR/Cas9 screens | % Dependent Cancer Cell Lines in RNAi screens |
| --- | --- | --- |
| PALB2 | 317/1150 = 27.5% | 5/550 = 0.9% |
| BRCA1 | 509/1150 = 44.2% | 16/710 = 2.3% |

We also directly compared CRISPRi to RNAi gene knockdown for many hits from the screen and found overall more efficient gene knockdown/larger phenotypes using CRISPRi, similar to the initial publication describing CRISPRi where they made the same comparison (Gilbert et al, 2014^27^, Fig. 1E). Consistent with our experience, the more limited set of ES cell lines in the RNAi screening database did not show a statistically significant difference in terms of PALB2 or BRCA1 dependence. Our results highlight important differences between genetic screening modalities (as noted previously by many groups^28,75,76^) and a potential advantage of CRISPR interference-based approaches for studying DNA repair genes.

## Supplemental Note 2

To quantify the extent of ATM suppression by EWSR1-FLI1, we benchmarked our effect size to loss of well-characterized activators of ATM including the MRN (MRE11/RAD50/NBS1) complex, MDC1, RNF8 and RNF168^41^ as described in the main text. As noted, there is a hierarchy amongst genes regulating ATM activity in response to DSBs: the MRN complex is required for ATM activation^47,48^ and serves as an upper bound for effect size, while the scaffolding protein MDC1^49,50^ and E3 ubiquitin ligases RNF8^51–53^ and RNF168^54,55^ have an intermediate phenotype given their important but not essential role in amplifying ATM signaling. Importantly, even loss of intermediate phenotype genes (i.e., MDC1^49^, RNF8^51^ or RNF168^54^) still produces a significant functional impact in terms of increased sensitivity to DSBs.

We utilized established MEF models of ataxia telangiectasia-like disease (ATLD)^45^ or Nijmegen breakage syndrome (NBS)^77^; human diseases of the MRN complex defined by genomic instability and radiosensitivity and caused by biallelic loss of function mutations in MRE11 or NBS1 respectively. These models allowed us to compare ATM signaling upon EWSR1-FLI1 expression with genetic loss of the MRN complex. As described previously, MRE11^ATLD/ATLD^ and NBS1^ΔB/ΔB^ MEFs display significantly decreased ATM autophosphorylation and minimal downstream ATM signaling (p-KAP1/p-CHK2) after IR compared to wild-type controls (Figs. S7A, B), with NBS1^ΔB/ΔB^ MEFs displaying a relatively more severe ATM phenotype than MRE11^ATLD/ATLD^ MEFs. EWSR1-FLI1 expression in wild-type MEFs suppressed IR-dependent ATM activation and downstream KAP1 and CHK2 phosphorylation to levels that are 38-56% of the effect size seen with MRE11^ATLD/ATLD^ or NBS1^ΔB/ΔB^ MEFs (Figs. S7A, B, E). We note that unlike MRE11^ATLD/ATLD^ or NBS1^ΔB/ΔB^ MEFs, the kinetics of ATM activation and CHK2 phosphorylation (as measured by mobility shift^45^, see Methods) are not delayed in EWSR1-FLI1 expressing MEF cells, consistent with residual ATM activity.

Additionally, we compared EWSR1-FLI1 expression to the milder ATM phenotypes induced by MDC1 or RNF168 shRNA knockdown in MEFs and observed overall similar levels of suppression in terms of ATM activation and signaling (Figs. S7C-G). There were differences between the trends within the two comparisons; EWSR1-FLI1 expressing MEFs displayed greater suppression of all three markers (p-ATM/p-KAP1/p-CHK2) when compared to RNF168 knockdown, whereas only p-KAP1 was decreased to a greater extent than MDC1 knockdown (though all three markers were still reduced relative to control MEFs). These data reinforce EWSR1-FLI1’s significant suppressive effect on IR-dependent ATM activation and signaling with an effect size comparable to knockdown of MDC1 or RNF168, but not equivalent (38-56%) to the near complete functional ATM deficiency seen with loss of the MRN complex in ATLD or NBS.

To assess the extent of ATM suppression at endogenous EWSR1-FLI1 levels in ES cells, we utilized the A673 “EWSR1-FLI1 degron” system. We compared auxin-induced degradation (AID) of endogenous EWSR1-FLI1 with shRNA knockdown of MRE11, NBS1, MDC1, RNF8, and RNF168. Consistent with the results in MEFs, the presence of EWSR1-FLI1 at endogenous levels suppressed IR-dependent ATM activation and downstream signaling to 29-64% of the levels seen with knockdown of MRN complex genes (Figs. S7H-K, R). To make these comparisons, we note that maximal ATM activity after IR was seen in the “shSCR (Scramble)/+Auxin” condition with no MRE11/NBS1 gene knockdown and EWSR1-FLI1 depleted for 3 hours prior to IR. To benchmark the ATM effects of endogenous EWSR1-FLI1 against MRN complex gene knockdown, we compared reduction from the maximal ATM activity condition (red bars) for (1) “shSCR/no Auxin (blue bars)” where EWSR1-FLI1 is now expressed at endogenous levels (and no MRN gene knockdown) versus (2) “shMRE11 or shNBS1/+Auxin (green bars)” where there is MRN gene knockdown (but EWSR1-FLI1 remains depleted).

In comparisons with MDC1 and the E3 ubiquitin ligases RNF8 and RNF168, endogenous levels of EWSR1-FLI1 suppressed IR-dependent ATM activation and signaling to an overall similar extent to loss of MDC1, RNF8, or RNF168 (Figs. S7L-R). There was again variation between the markers with endogenous EWSR1-FLI1 having comparable (or slightly greater) levels of p-ATM and p-KAP1 suppression compared to MDC1, RNF8 or RNF168 knockdown, whereas knockdown of MDC1 (but not RNF8 or RNF168) had a greater effect on CHK2 phosphorylation consistent with a direct interaction between MDC1 and CHK2^78^. Across all experiments, the combined presence of endogenous EWSR1-FLI1 (no Auxin) and gene knockdown could further suppress ATM activation/signaling even in MRE11 or NBS1 knockdown cells, suggesting non-redundant mechanisms of ATM suppression. Taken together, our results demonstrate that EWSR1-FLI1 suppresses IR-dependent ATM activation and signaling with an effect size that is not equivalent (∼30-60%) to the near complete absence of ATM activity seen with loss of the MRN complex, but comparable to loss of classical ATM activators including MDC1, RNF8, and RNF168 (genes whose loss still confer significant DSB sensitivity) (Figs. 5G, R). In summary, these data contextualize the effect size and biological significance of the partial ATM defects in ES.

## Supplementary Note 3

We note here the significant disparity between the relatively rare occurrence of TP53 (6-10%) mutations in ES patient tumors^34,79^, especially at diagnosis, compared to the frequent TP53 mutations seen in ES cell lines. For example, 13 of the 17 annotated ES cell lines in DepMap are TP53 mutant. There is a similar but less dramatic discrepancy between the frequency of both STAG2 mutations (15-20%) and CDKN2A/B loss (12-15%) in patient tumors compared to ES cell lines. We therefore included cell lines with wild-type TP53, STAG2, and CDKN2A/B (none are wild-type for all three to our knowledge) to address possible contributions to the observed ATM effects. The TP53, STAG2, and CDKN2A/B status of the major cell lines used in this manuscript are:

1. A673 (TP53 mutant, STAG2 wild-type, CDKN2A/B loss)^34,80^
2. TC-32 (TP53 wild-type, STAG2 mutant, CDKN2A/B loss)^81,82^
3. TC-252 (TP53 wild-type, STAG2 mutant, CDKN2A/B loss)^82,83^
4. ES-8 (TP53 mutant, STAG2 mutant, CDKN2A intact)^13,82^

Thus, we include data from 2 cell lines with wild-type TP53 (TC-32 and TC-252), 1 cell line with wildtype STAG2 (A673) and 1 cell line with the CDKN2A/B locus intact (ES-8). We also validate the core findings of ATM suppression and ATR inhibitor sensitivity using an inducible EWSR1-FLI1 shRNA in A673 (STAG2 wild-type) and TC-32 (TP53 wild-type) cells (Fig. 4). Therefore, we conclude that the observed ATM effects do not depend on TP53 (or STAG2, CDKN2A/B) status, though we do not fully exclude possible contributions. We also note that TP53 and STAG2 status have important downstream consequences for cell viability^84^, metastatic potential^80^, and clinical outcomes in patients^34,79^ as previously described.

## Methods

### Cell lines and culture conditions

U2OS, DL-221, HeLa, and 293T cells were cultured in Dulbecco’s Modified Eagle Medium (DMEM)-High glucose (Cytiva) supplemented with 10% fetal calf serum (FCS) and streptomycin/penicillin (100 μg/ml). A673, TC-32, ES-8, RD-ES, SU-CCS-1, A549, H3122, and HCC364 cells were grown in RPMI-1640 (Gibco) supplemented with 10% FCS and streptomycin/penicillin (100 μg/ml). BER and BOD cells were cultured in DMEM/F12-High glucose with 10% FCS. Cell lines were maintained at 37 °Celsius in a humidified incubator with 5% CO2. All cell lines were subjected to STR analysis and mycoplasma testing.

A673 cells with a doxycycline(dox)-inducible shRNA against EWSR1-FLI1 were a kind gift from Franck Tirode^31^ (clone 1c); the shRNA sequence (GGCAGCAGAACCCTTCTTAT) targets the EWSR1-FLI1 junction. TC-32 cells with a dox-inducible shRNAs against EWSR1-FLI were a kind gift from Dr. David McFadden and the shRNA sequence (ATCCGACCGAGTCGTCCATGTA) targets the 3’ portion of FLI present in the EWSR1-FLI fusion (native FLI1 is not expressed in TC-32 cells). A673 “EWSR1-FLI1 degron” cells were also a kind gift from Dr. David McFadden and the experimental details regarding cell line generation are referenced in McGinnis et al, 2024, bioRxiv^44^. MEF models of ataxia telangiectasia-like disease (ATLD)^45^ or Nijmegen breakage syndrome (NBS)^77^ were a kind gift from Dr. John Petrini.

### CRISPR-interference screen

Ewing sarcoma cell line A673 was transduced with dCas9-KRAB-BFP and then doubly sorted for BFP positive cells. Lentiviral particles were generated from 293T cells transduced with pooled sgRNA libraries (7000 genes, 10 guides per gene) as described previously^27^ and then used to infect A673 dCas9-KRAB cells at a multiplicity of infection of 0.3. Cells were selected in puromycin for 48 hours and an aliquot was frozen for t_0_ analysis. The remainder of cells were seeded in 500 cm^2^ tissue culture plates at equal density (1 × 10^7^ cells/plate) and then treated with vehicle (DMSO) or 12.5 nM/40 nM doxorubicin for 72 hours. After 72 hours, cells were trypsinized, counted and pooled every 3 days, and then replated in media without doxorubicin at the same density to maintain minimum 500× coverage of each sgRNA construct. Cells were viably frozen after 10 days and cell counts were used to determine actual lethal dose (LD) values. 2 biologic replicates were performed, and data combined.

Deep sequencing and data analysis were performed as described^27^. Briefly, genomic DNA was extracted from t_0_ and t_end_ cells using a DNEasy Blood & Tissue kit (Qiagen), digested to enrich for lentiviral integration sites, and sgRNA sequences were amplified by PCR for subsequent sequencing on an Illumina HiSeq. Reads were aligned to the sgRNA library and fold-change from t_0_ to t_end_ in the DMSO, low and high doxorubicin conditions were calculated. A gene-level score was then calculated as the mean of the top 3 scoring sgRNAs targeting a given transcript.

### CRISPRi growth assays

dCas9-KRAB expressing cells were infected with gRNAs (sequences below) and then selected with puromycin for 48 hours. 50,000 cells were then re-plated in a 6 well plate and cell counts were obtained by trypsinization and counting at day 5. For experiments in A673 cells with dox-inducible shRNA against EWSR1-FLI1, 1μg/ml Dox (or vehicle) was added upon replating the puromycin selected, gRNA-expressing cells. gRNA sequences are as follows:

sgBRCA1-1: GCGTAAGGCGTTGTGAACCCT

sgBRCA1-2: GCTCGCTGAGACTTCCTGGAC

sgPALB2-1: GCGCACTGAGGGTGCGATCC

sgPALB2-2: GATTTAATTGGCCGGAGTTT

sgControl-1: GGCTCGGTCCCGCGTCGTCG

sgControl-2: GAACGACTAGTTAGGCGTGTA

### Computational analysis of genomic data

Whole genome sequencing (WGS) data from known BRCA mutant/wildtype breast cancer patient samples and Ewing sarcoma samples were obtained from the European Genome-phenome Archive (EGA). The WGS data for the ES cohort was published in Tirode et al, 2014^34^ and deposited with study accession numbers: EGAS00001000855 (Institute Curie cohort) and EGAS00001000839 (St. Jude’s cohort). The breast cancer WGS was reported in 2016^35^ and deposited under EGAD00001001322, as part of a larger WGS of 560 breast cancer samples (overall project accession number EGAS00001001178).

FASTQ files were aligned to the GRCh38 (patch 5) human genome reference using BWA MEM^85^, with pre- and post-alignment filtering and processing using NGSUtils^86^ and GATK^87^. Insertions and deletions (indel) were called using MuTect 2 for short indels^88^ and Delly for larger indels^89^. For each indel, the amount of microhomology at the flanking ends was calculated using the custom software “mhscan”. Briefly, for each indel, mhscan will determine the number of bases of homology between the insert or deletion and the flanking sequence. All indels with between 2-6 bp of homology were classified as microhomology (MH) positive. Mutational signatures were calculated using the somatic single-nucleotide variants for each sample. COSMIC v2^90,91^ signatures were calculated using deconstructSigs^92^.

### Determining deletion length bins

The binning ranges were determined by calculating the ability for the number of deletions of each length (1-100 bp) to accurately separate the BRCA wild type (wt) / mutant (mut) samples into two groups. The deletions were classified as either MH-positive or MH-negative and the total number of deletions for each length was calculated for BRCA wt and BRCA mut samples (data were aggregated for all samples in the group). For each deletion length, a Fisher exact test was used to calculate the statistical significance of whether the MH-positive to MH-negative ratio separated the BRCA wt/mut groups (Fig. S2F). The distribution of p-values was used to determine appropriate bin sizes for downstream analysis. Deletions were binned into ranges of 1-6 bp, 7-28 bp, 29-45 bp, 46-100 bp, and 101-1000 bp.

### DSB Repair reporter assays

HR, Alt-EJ, SSA and c-NHEJ efficiencies were measured using previously established DR-GFP, EJ2-GFP, SA-GFP and EJ5-GFP reporter systems respectively^38^. Briefly, dual promoter plasmids co-expressing mTagBFP and EWSR1-FLI1 (or empty vector, EV) were generated by replacing the puromycin cassette with mTagBFP in the plasmid pCDH-CMV-EWSR1-FLI1 (Addgene #102813). Co-expression plasmids were delivered by nucleofection into U2OS DSB reporter cell lines using the SE Cell Line 4D-Nucleofector XL Kit (Lonza Biosciences). To induce a DSB, the I-SceI-P2A-mCherry plasmid which we generated by replacing AmCyan with I-SceI in the bicistronic plasmid Amcyan-P2A-mCherry (Addgene #45350) was delivered by nucleofection after 24 hours. Flow cytometry was performed 72 hours after I-SceI introduction. DSB repair efficiency was calculated by determining the percentage of GFP positive cells within the BFP and mCherry double-positive population (see Fig. S3A). For siRNA experiments, siRNA (5nM) was delivered by transfection using Dharmafect transfection reagent (Dharmacon) 72 hours prior to I-SceI introduction. To directly examine junctional sequences after DSB induction for evidence of cNHEJ or alt-EJ (Fig. S3B), we utilized a distinct I-SceI based DSB repair reporter described here^40^. Briefly, 5μg of the NHEJ reporter was transfected into EWSR1-FLI1 (or EV) expressing U2OS cells, GFP+ cells were sorted at 72 hours for genomic DNA isolation, and then DSB junctional sequences analyzed by recircularization of the repaired NHEJ reporter and plasmid sequencing.

### DDR signaling experiments

For experiments in A673 and TC-32 cells with dox-inducible shRNA against EWSR1-FLI1, knockdown was induced by either addition of 1μg/ml dox (A673) or 200 ng/mL (TC-32) for 72 hours followed by IR treatment. For A673 “EWSR1-FLI1 degron” cells, 200 μM IAA (auxin, Sigma) was added 3 hours prior to IR treatment for EWSR1-FLI1 depletion. For the ectopic expression experiments in U2OS cells, the respective fusion proteins (EWSR1-FLI1-GFP, EWSR1-ATF1-GFP, or EV-GFP) were introduced via nucleofection with homemade buffers (Solution 1: 2g ATP-disodium salt, 1.2g MgCl_2_.6H_2_0 in 10ml water; Solution 2: 6g KH_2_PO_4_, 0.6g NaHCO_3_, 0.2g Glucose in 500ml water, pH 7.4. Solutions were stored at -20°C and 80μl of Solution1 was mixed with 4ml of solution 2 at the time of nucleofection). GFP positive cells were selected by FACS and 24 hours after sorting, subjected to IR. MEF cells were infected with pCDH-EWSR1-FLI1 (Addgene plasmid #102813) lentivirus or EV, selected with puromycin within 72 hours, and then subjected to IR.

For the shRNA experiments testing knockdown of ATM regulators in A673 “EWSR1-FLI1 degron” or MEF cells, Mission pLKO.1 shRNA viruses (sequences below) were used to infect target cells and selection with puromycin was completed within 72 hours to minimize effects on cell viability, followed by IR treatment. For experiments in SU-CCS-1 cells, siRNA against EWSR1-ATF1 was delivered by nucleofection 72 hours prior to IR. For improved induction of ATR/CHK1 signaling, cells were treated with neocarzinostatin (NCS) at 200ng/mL for 30 minutes. Lysates were collected at defined time points post-IR (or post-NCS) and subjected to Western blotting.

### Western blotting and quantification

Cells were washed with 1X PBS and scraped in RIPA buffer (25 mM Tris⋅HCl (pH 7.6), 150 mM NaCl, 1% NP-40, 1% sodium deoxycholate, 0.1% SDS), supplemented with 1X HALT protease inhibitor cocktail and 1X HALT phosphatase inhibitor cocktail. Cells were lysed using a syringe and cellular debris were separated by centrifugation. Lysates were quantified using Bradford’s reagent and 15μg lysate was loaded onto SDS-PAGE gels followed by blotting of separated proteins. Blots were blocked with 5% BSA in TBS-T for 30 minutes at room temperature followed by incubation with primary antibody overnight at 4°C. After 1 hour incubation with the appropriate HRP-conjugated secondary antibody, signal was detected using ECL Prime reagent (Amersham, Cytiva) on an ImageQuant LAS 4000. Quantifications were performed using Fiji and all protein levels were normalized to loading control Actin. To measure CHK2 phosphorylation in MEFs as no reliable mouse p-CHK2 antibody is commercially available, we quantified the intensity of the upper band (after IR) on total CHK2 immunoblots which corresponds to the phosphorylated species, as utilized previously by multiple groups^46,49^.

The list of antibodies utilized in this study includes: FLI1 (Abcam, ab133485, 1:500), CtIP (Cell Signaling Technologies (CST), D76F7, 1:000), 53BP1 (Abcam, ab153909, 1:1000), p-ATM (Invitrogen, MA5-32751), ATM (CST, 2873S, 1:1000), pCHK2-T68 (CST, 2661S, 1:1000, Human only), CHK2 (CST, 2662S, 1:1000), p-KAP1 (Novus Biologicals, A700-013, 1:1000), KAP1 (Novus Biologicals, A700-014, 1:1000), p-DNA-PK (CST, 68716S, 1:1000), DNA-PK (CST, 4602T, 1:1000), p-ATR (Genetex, GTX128145, 1:1000), ATR (Novus Biologicals, A300-138A, 1:1000), pCHK1-Ser317: (CST, 2344, 1:1000), CHK1 (CST, 37010, 1:1000), GFP (CST, 2956S, 1:1000), EWSR1 (Genetex, GTX114069, 1:1000), beta-Actin (Sigma, A2228, 1:2000), anti-Flag (Sigma, F3165, 1:2000), ATF1 (Novus Biologicals, A303-036A, 1:1000), FUS (Novus Biologicals, A300-302A-1:1000), RNaseH1 (Genetex, GTX117624, 1:1000), MRE11 (CST, 1:1000, 4895S), NBS1 (CST, 1:1000, 14956), RAD50 (CST, 1:1000, 3427S), MDC1 (Invitrogen, MA5-27650, 1:1000), RNF8 (Proteintech, 1:1000, 14112-1-AP), RNF168 (EMD Millipore, 1:1000, ABE367, Mouse only), and RNF168 (Thermo, 1:1000, H00165918-MO1, human).

### siRNA/shRNA sequences

#### siRNA sequences

EWSR1: Mix of two siRNAs targeting the C-terminus of EWSR1: GGAACAGAUGGGAGGAAGA and AGGAAAGCCCAAAGGCGAU

CTIP: On-TARGETplus SMARTpool siRNA (Dharmacon, L-011376-00-0005)

BRCA1: On-TARGETplus SMARTpool siRNA (Dharmacon, L-003461-00-0005)

POLQ: On-TARGET plus SMARTpool siRNA (Dharmacon, L-015180-01-0005)

53BP1: On-TARGETplus SMARTpool siRNA (Dharmacon, L-003548-00-0005)

MRE11: On-TARGETplus SMARTpool siRNA (Dharmacon, L-009271-00-0005)

EWSR1-ATF1: GCGGUGGAAUGGGAAAAAUUU

#### shRNA targeting sequences (pLKO.1 Mission shRNA)

LIG4: TATGTCAGTGGACTAATGGAT

Mouse MDC1: mixture of 2 shRNAs, AGCATGCCTCACTCCTATAAG and GAGCCTCAATGGCACTCAAAT

Mouse RNF168: mixture of 2 shRNAs, GCCAACTTCTACTCAAGATAA and CCTTGGCTTCTCCTTTGAGTT

Human MRE11A: mixture of 2 shRNAs, ACGACTGCGAGTGGACTATAG and TGTTGGTTTGCTGCGTATTAA

Human NBS1: mixture of 2 shRNAs, CCTCTTGATGAACCATCTATT, and GCTCGAAAGAATACAGAACTA

Human MDC1: mixture of 2 shRNAs, CCCTGAATCAACTGTCCCTAT, CGGACCAAACTTAACCAAGAA

Human RNF168: mixture of 2 shRNAs, GCAGTCAGTTAATAGAAGAAA, and CGTGGAACTGTGGACGATAAT

Human RNF8: mixture of 2 shRNAs, TGGAGCAACTAGAGAAGACTT, and CCAAAGAATGACCAAATGATA

### Cell survival assays

350,000 cells were seeded in 6-well plates, 24 hours prior to drug treatment. Cells were treated with varying doses of ATR kinase inhibitor elimusertib (Selleckchem) +/- doxorubicin (Sigma) or CHK1 inhibitor LY2603618 (Selleckchem) for 3 days at the end of which cells were counted to determine fractional cell survival. For dox-inducible EWSR1-FLI1 knockdown in A673 and TC-32 cells, shRNA was induced by treatment with dox for 72 hours prior to ATR or CHK1 inhibitor treatment. For RNAseH1 experiments, A673 cells were infected with pLV-EF1a-RNAseH1-IRES-Blast lentivirus (derived from Addgene plasmid #85133) or empty vector and selected for stably infected cells.

### Laser micro-irradiation

Laser micro-irradiation was performed as previously described^21^. Briefly, U2OS cells expressing EWSR1-GFP, EWSR1-FLI1-GFP, EWSR1-ATF1-GFP, EWSR1-WT1-GFP or the various mutant forms of the fusion oncoproteins were seeded in 8-well Lab Tek II Chamber Slides (Thermo Fisher Scientific) for 24 hours. Cells were treated with 1µg/ml Hoechst 33342 (Thermo Fisher Scientific) for 30 minutes prior to micro-irradiation. Live cell microscopy was performed using Nikon Ti microscope with a CSU-W1 spinning disk confocal using a 100X/1.4 Plan Apo VC objective at the UCSF Center for Light Microscopy. To induce DNA damage, 5-pixel wide stripes were drawn in every cell nucleus to label the region of interest (ROI) and irradiated with a 405nm diode laser (40mW). Images were acquired pre-irradiation and at 1-minute intervals post-laser damage for 15 minutes. To plot recruitment kinetics, the pre-irradiation fluorescence intensity was subtracted from the intensity of the ROI for every nucleus.

### Co-Immunoprecipitation (co-IP) experiments

For EWSR1 co-IP experiments, 293T cells were transfected with Flag-EWSR1 and EWSR1-FLI1 (or empty vector) for 48 hours. Cells were then treated with NCS (200ng/mL) for 30 minutes and cells were collected at 15 minutes, 30 minutes and 1 hour post NCS treatment. Nuclear co-IP was performed using the Nuclear Co-IP kit (Active Motif) according to the manufacturer’s instructions followed by IP using M2 Flag beads (Sigma). For GFP co-IP experiments, 293T cells were transfected with EWSR1-FLI1-GFP, FLI1-GFP, EWSR1-GFP, or GFP empty vector for 24 hours, followed by IR treatment (5Gy), cell collection at specified time points, the same nuclear co-IP protocol, and IP using ChromoTek GFP-Trap Agarose beads (Proteintech).

### Cell cycle analysis

U2OS cells were nucleofected with the dual promoter pCDH-EWSR1-FLI1-EF1a-mtagBFP (or empty vector) and analyzed at 48 hours. Cells were fixed using 4% formaldehyde for 15 minutes at room temperature. After washing with 1% BSA in PBS, cells were resuspended in 500μl of FxCycle™ PI/RNAse Solution (ThermoFisher Scientific) for 30 minutes at room temperature. Cell cycle profiles were also analyzed using Click-iT EdU Alexa Fluor 488 Flow Cytometry Assay Kit (Invitrogen). Cells were pulse labelled with 10μM EdU for 30 minutes and processed according to the manufacturer’s instructions. Flow cytometry was used to analyze the cell cycle profile for BFP positive, EWSR1-FLI1 (or empty vector) expressing cells.

### Patient-derived xenograft (PDX) experiments

The establishment of PDX models was performed by the laboratory of Dr. Filemon Dela Cruz and Dr. Andrew Kung from patients at MSKCC according to the institutional animal and IRB protocols. Tumor fragments from patients were serially transplanted into athymic nude (Foxn1^nu^) mice from Charles River Laboratories to establish the PDX for 3-5 passages, after which time seeds were implanted to begin the PDX experiments. When average tumor volumes exceeded 150mm^3^, mice were randomized into either elimusertib or vehicle group. Elimusertib was dosed at 40 mg/kg twice daily, 3 days-on/4 days-off as previously reported^71^, and dissolved in 60% polyethylene glycol 400, 10% ethanol, and 30% water to a 4 mg/mL solution. The same solution without compound was used as vehicle control. Mice were sacrificed by cervical dislocation once the tumor volume exceeded 2000 mm^3^ or body weight loss was higher than 20% (which did not occur), and tumors were collected for immunohistochemistry (IHC).

For IHC studies, paraffin cross-sections (thickness = 5μm) of PDX tumor tissues were applied to Superfrost Plus microscope slides (Fisher, #1255015) for immunohistochemical analyses. After heat-mediated antigen retrieval (48 minutes, Cell Conditioning 1) (Ventana, #950-500), PDX tumor cross-sections were incubated for 4 hours at room temperature with the following primary antibodies: rabbit polyclonal Cleaved Caspase-3 (Cell Signaling Technology, #9661, 0.05mg/mL), rabbit monoclonal Ki67 (Cell Signaling Technology, #9027, 0.25mg/mL), gH2AX Ser139 (Cell Signaling Technology, #9718, 1:1,000 dilution), p-CHK1 (Cell Signaling Technology, #2348, 1:100 dilution), or normal rabbit polyclonal control IgG (Invitrogen, #10500C). After sequential washes with PBS, sections were incubated for 20 minutes with Omni Map anti-Rb HRP (Roche, #760-4311), followed by DAB ChromoMap Kit amplification (Ventana Medical Systems, #760-159). The slides were counter-stained with hematoxylin and mounted with Permount (Fisher Scientific). Analysis was performed in a blinded fashion using QuPath software.

### Statistics

Data from at least three independent experiments were used to calculate P values which were determined with Student’s t tests and ANOVA using GraphPad software.

## Data and Code Availability

All data supporting the findings of this study are available within the paper and its Supplementary Information. Code for “mhscan” is available on request.

## Author Contributions

SM, DG, TGB, ASC, and AT conceived and designed the study. SM, DG, YPL, AFG, HA, SP, NW, AH, and AT conducted the experiments and collected data. MB designed the computational algorithms and analyzed data. FDC, TF, and EDS performed the PDX experiments. SM, DG, MH, TGB, and AT analyzed and interpreted the data. JW and ASC provided support for the CRISPR screening and computational analysis respectively. SM, TGB, and AT wrote the manuscript.

## Acknowledgements

This project is supported by the Tow Center for Developmental Oncology, the V Foundation, Hyundai Hope on Wheels, the Sarcoma Center at MSKCC, NIH/NCI Cancer Center Support Grant P30 CA008748, Alex’s Lemonade Stand, the A.P. Giannini Foundation, and the St. Baldrick’s Foundation (to A.T), as well as NIH/NCI U54CA224081, R01CA169338, R01CA211052, R01CA204302, U01CA217882 (to T.G.B). A.T. was an advisor to Faze Medicines. T.G.B. is an advisor to Novartis, Astrazeneca, Revolution Medicines, Array/Pfizer, Springworks, Strategia, Relay, Jazz, Rain, EcoR1 and receives research funding from Novartis and Revolution Medicines and Strategia. The authors would like to acknowledge A. Yasemin Goksenin and Zoji Bomya for experimental help, Amit Sabnis, Alan Ashworth, John Petrini, and Alex Kentsis for scientific input and manuscript review, Jeremy Stark for the DNA double-strand break reporter cell lines, the Center for Advanced Light Microscopy at UCSF for technical help, and the Molecular Cytology Core Facility at MSKCC for IHC staining. We especially thank the lab of Dr. David McFadden for sharing unpublished reagents including the A673 “EWSR1-FLI1 degron” cells and TC-32 cells with dox-inducible shRNAs against EWSR1-FLI, as detailed in the Methods.

